# Dual credit assignment processes underlie dopamine signals in a complex spatial environment

**DOI:** 10.1101/2023.02.15.528738

**Authors:** Timothy A. Krausz, Alison E. Comrie, Loren M. Frank, Nathaniel D. Daw, Joshua D. Berke

**Affiliations:** Neuroscience Graduate Program, University of California, San Francisco; Kavli Institute for Fundamental Neuroscience, and Weill Institute for Neurosciences, UCSF; Howard Hughes Medical Institute; Department of Physiology, UCSF; Department of Psychology, and Princeton Neuroscience Institute, Princeton University, NJ; Department of Neurology, and Department of Psychiatry and Behavioral Science, UCSF

**Keywords:** Dopamine, Credit Assignment, Reinforcement Learning, Maze

## Abstract

Dopamine in the nucleus accumbens helps motivate behavior based on expectations of future reward (“values”). These values need to be updated by experience: after receiving reward, the choices that led to reward should be assigned greater value. There are multiple theoretical proposals for how this credit assignment could be achieved, but the specific algorithms that generate updated dopamine signals remain uncertain. We monitored accumbens dopamine as freely behaving rats foraged for rewards in a complex, changing environment. We observed brief pulses of dopamine both when rats received reward (scaling with prediction error), and when they encountered novel path opportunities. Furthermore, dopamine ramped up as rats ran towards reward ports, in proportion to the value at each location. By examining the evolution of these dopamine place-value signals, we found evidence for two distinct update processes: progressive propagation along taken paths, as in temporal-difference learning, and inference of value throughout the maze, using internal models. Our results demonstrate that within rich, naturalistic environments dopamine conveys place values that are updated via multiple, complementary learning algorithms.

## Introduction

Animals frequently make motivated choices based on prior experiences - for example, selecting paths towards locations where food was previously found. Achieving such adaptive decision-making can pose a computational challenge. In particular, decision points can be separated from rewards by considerable gaps in time and space. When rewards are obtained (or unexpectedly omitted) this separation produces a “credit assignment problem”: determining which places and choices ought to gain or lose value. The specific algorithms that brains use to solve this problem are not well understood.

Reinforcement Learning (RL) theory provides an array of candidate algorithms for generating and updating value signals (1). In “temporal difference” (TD) learning, value is passed between sequentially experienced states (situations). In brief, as each state is encountered its associated value becomes eligible for updating. Unexpected rewards, or transitions to states with unexpected values, evoke “reward prediction errors” (RPEs). RPEs are learning signals: they update the values of eligible states. In this way, values can be progressively propagated back to earlier states, over repeated episodes of experience. Temporal difference RPEs can be encoded by brief (phasic) changes in the firing of midbrain dopamine (DA) cells (2–5), and by corresponding changes in DA release in the nucleus accumbens (NAc; (5, 6)). However, despite the compelling correspondence between phasic DA and TD RPEs, current evidence that value propagates along sequences of states in a TD-like manner is sparse at best (7–9).

TD learning is a “model-free” (MF) algorithm: learning occurs only from direct experience of states, without using knowledge of how those states are related. A complementary set of “model-based” (MB) algorithms can achieve greater flexibility in learning and decision-making by using knowledge about state relationships to infer and update values. For example, after taking one path and receiving reward, MB algorithms can increase values along *alternative* paths to the same reward location (10, 11). In at least some behavioral contexts, DA signals reflect RPEs that incorporate such inferred information (12–15).

NAc DA release also gradually ramps up as animals actively approach expected rewards (5, 16–19). These ramps appear to signal the value of the upcoming reward, discounted by spatial distance (although they have also been interpreted as RPEs (20, 21)). As DA ramps are more apparent when the behavioral context favors use of an internal model (22), they have been proposed to reflect ongoing MB calculations.

Yet overall, existing evidence does not tease apart the specific algorithms used to estimate and update values, or reveal how these values are reflected in DA signals. Many behavioral tasks (e.g. (5, 23, 24)) commonly used to investigate DA and value coding involve only minimal separation between an action and its outcome, thus avoiding the challenging credit-assignment question. In other paradigms, applying RL ideas involves unsupported arbitrary assumptions (25) – e.g., choosing the set of discrete covert states to span a time interval between a cue and reward (2). Spatial tasks have the advantage that the brain has a well-studied set of spatial representations that could serve as a basis for RL states (26). However, most spatial tasks – especially those in which DA dynamics have been investigated – are very simple (e.g., T-mazes;(19, 27)). This simplicity is often useful, but can prevent critical tests that distinguish between credit assignment algorithms.

To better elucidate MF and MB credit-assignment processes within natural environments, we developed a dynamic, complex spatial foraging task for rats, the Hex Maze. In this task, animals traverse through numerous distinct decision points in the pursuit of reward, and choices are separated from their outcomes by multiple steps in space and time. Furthermore, reward contingencies can be unstable, and the available paths to reward locations can be unexpectedly reconfigured. We show that rats readily adapt to these changes, and incorporate both costs (current distances to reward ports) and benefits (current reward probabilities) into their decisions.

Using fiber photometry, we observe DA RPE coding at reward receipt and also strong DA pulses when rats discover newly available paths. We confirm that NAc DA ramps up with reward approach, and we show that these ramps reflect a robust relationship between DA release at each location and the dynamically changing value of that location. We then take advantage of this DA place-value signal to examine how values are updated from trial-to-trial. We report strong evidence for *both* MF TD-like local propagation of values between adjacent locations, and MB global inference of values throughout the maze environment.

## Results

### Cost-benefit decision-making in the Hex Maze

The Hex Maze (Fig. 1; Supp. Video 1) contains a reward port at each corner, each with a distinct reward probability (17, 28–31). The available paths to these reward ports are defined by a set of barriers, constraining rats into making sequences of left and right turns from each “hex” location. The task is self-paced- the end location for each “trial” is the start for the next - and each reward port can be approached from multiple starting locations. Overall, rats (n=10) were more likely to choose a port if it had a higher probability of reward (Fig. 1b), and was closer (Fig. 1c), compared to the alternative. A mixed-effects logistic multiple regression, incorporating any turn biases (Methods), revealed highly significant effects of both reward probability (mean *β* = 1.613 ± 0.158 SEM, p = 1.91 * 10^−24^) and distance cost (mean *β* = -6.803 ± 0.549 SEM, p = 2.61 * 10^−35^) on port choices. After each block of 50-70 trials (traversals between ports), either the reward probabilities changed (Fig. 1e) or a barrier is moved to change available paths (Fig. 1f). After a change in reward probabilities, rats increased their choice of ports whose reward probability has increased (Fig. 1g). Following a barrier move, rats adjusted their port choices to favor shorter paths (Fig. 1h) and also progressively refined their specific paths to be more efficient (Supp. Fig. 1).

**Fig. 1.**
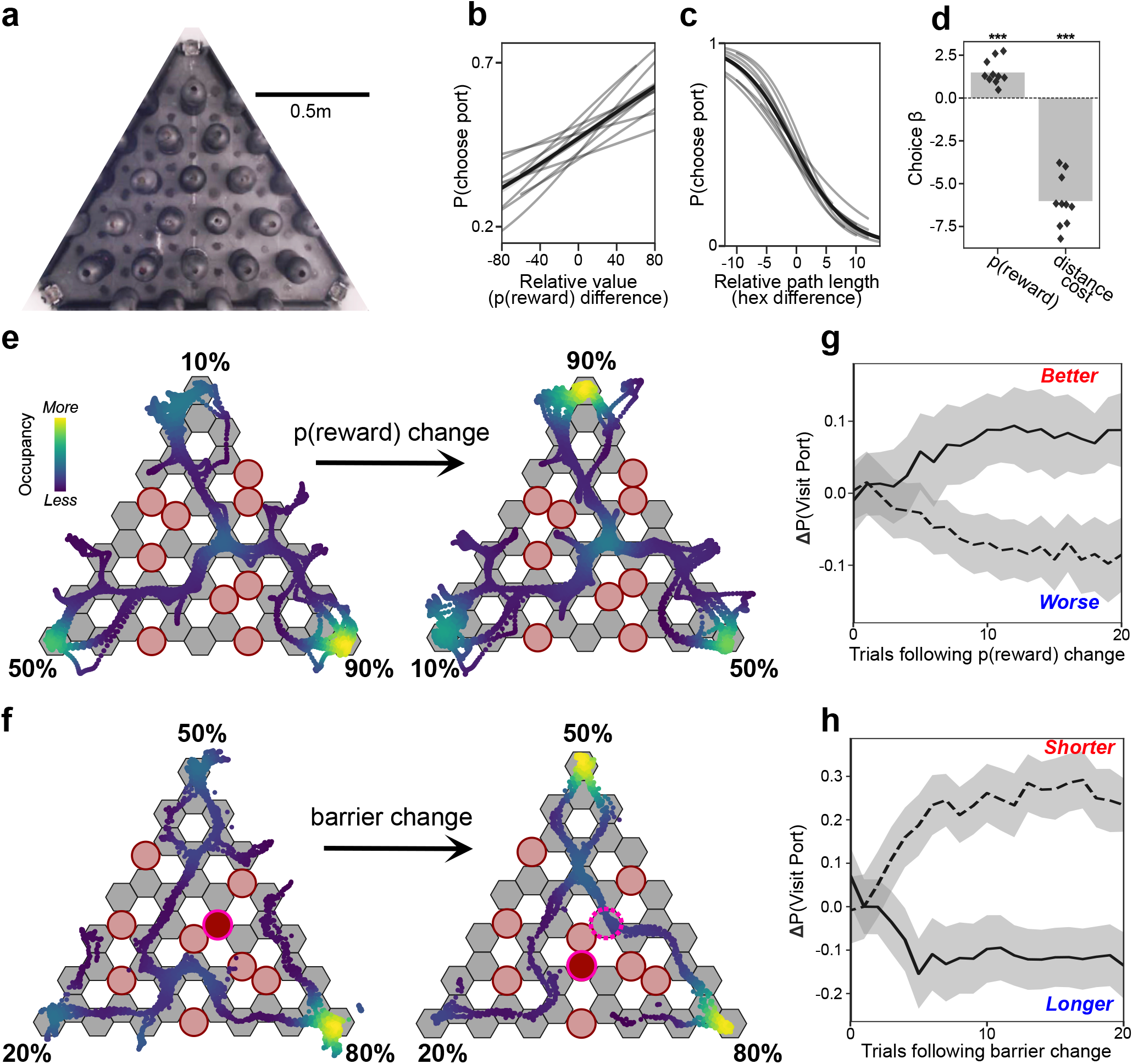
Adaptive behavior in the Hex Maze. **a**, Birds-eye view of the maze. Permanent barriers (black columns) divide the area into 49 hexagon-shaped choice points (“hexes”). Additional movable barriers (absent here) determine the available paths to the reward ports at each corner. Once visited, a port’s reward probability becomes zero until another port is visited. **b**, Probability of choosing an available port (leftward, from the current port) as a function of the difference between that port’s and the alternative port’s reward probabilities. Grey traces are individual-rat logistic curves fit to the data, and the black line shows the mean relationship. **c**, Same as b, but a function of the difference between path lengths to the available ports. **d**, Results of logistic multiple regressions run for each individual rat, showing the positive influence of reward probability and the negative influence of path length on choices. Significance asterisks are from the mixed-effects regression analysis. For b-d, only the second half (trial number > 25) of each block was included to allow rats time to adapt to changes (n = 10 rats, 82 sessions, 9079 trials). **e**, Example of a reward probability change. Red circles indicate hexes containing a movable barrier, dots show the rat’s detected positions (color coded by occupation density; second halves of blocks). White, empty, hexes indicate the positions of the permanent barriers shown in a. **f**, Example of a barrier change. Dark red circle with a pink outline shows the moved barrier. **g**, Mean change in port choice probability following increases (solid line) or decreases (dashed line) in reward probability (n = 10 rats, 36 sessions, 134 blocks; error bands indicate *±* SEM). “Trials” here include those where the rat had the opportunity to choose the port in question. **h**, Mean change in port choice probability following increases (solid line) or decreases (dashed line) in the path length to get there (n = the same 10 rats, 46 sessions, 162 blocks; error bands indicate *±* SEM).”Trials” here include those where the rat had the opportunity to choose the port in question.

### Phasic dopamine responses to rewards and novel path opportunities

During Hex Maze performance we recorded NAc DA dynamics using fiber photometry with the fluorescent DA sensor, dLight1.3b (32); (n = 10 rats, 19 fiber locations, 82 behavioral sessions, 296 blocks, 16379 trials, mean of 1638 trials per rat). We first examined DA changes around reward port entry, since receipt (and omission) of probabilistic reward is an obvious time to look for the best-known correlate of NAc DA, RPE signals. DA transiently increased or decreased depending on whether reward was delivered or omitted, respectively (Fig. 2b). The magnitude of these phasic changes depended on port reward probability, in a direction consistent with RPE coding (Fig. 2c, Pearson correlation, rewarded trials mean coefficient = -0.214 +/-0.99 STDEV; omission trials mean coefficient = -0.107 +/- 0.0.068 STDEV; both significantly different to zero across n=10 rats, two-tailed Wilcoxon Signed Rank tests, p = 1.95*x*10^−3^ each). To better estimate RPE at the single-trial level, we fit a simple trial-level RL algorithm to rats’ port choices and reward outcomes (“Q learning”; see Methods). DA following port entry significantly scaled with these RPE estimates (Supp. Fig. 2), although encoding of positive RPEs was notably stronger and more consistent across rats, compared to negative RPEs (in line with prior studies; (3, 5, 33)).

**Fig. 2.**
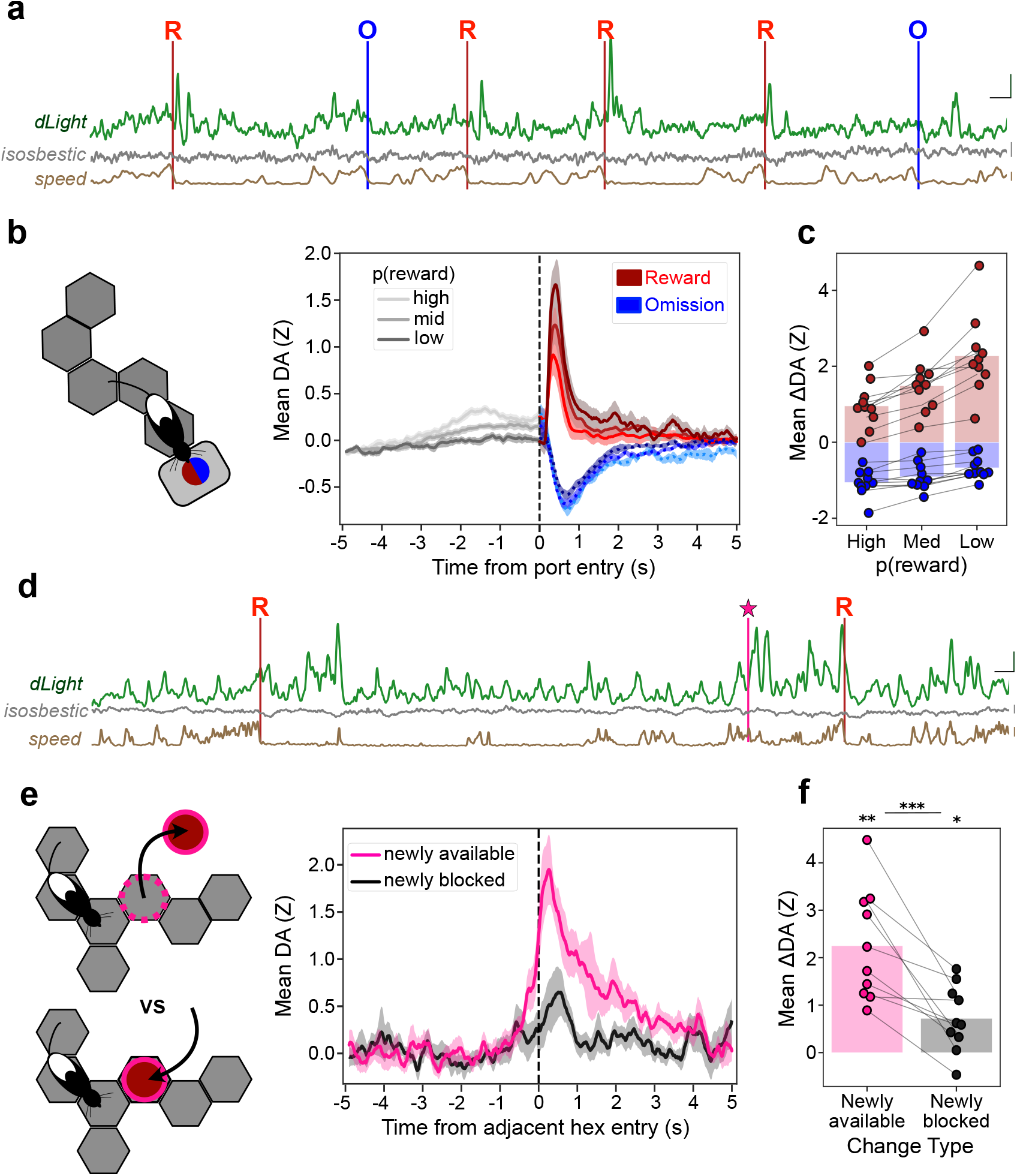
Dopamine pulses at rewards and novel path opportunities. **a**, Example trace of dLight, isosbestic (405nm) control signal, and running speed over three trials. Red “R”s and vertical lines indicate moments of reward delivery, and blue “O”s and vertical lines indicate reward omissions upon port entry. Vertical scale bars indicate 2Z for fluorescence signals and 20cm/s for speed. Horizontal scale bar indicates 2s of time. **b**, *Left*, cartoon of rat arriving at port. *Right*, average DA (Z-scored) aligned to port entry, pooled by the destination port’s reward probability (“high”, 80 or 90%; medium, 50%; “low”, 10% or 20%). Traces are separated into rewarded (red) or omissions (blue) following port entry, and error bands indicate *±* SEM (n=10 rats). Only the second half (trial number > 25) of each block was included (82 sessions, 9079 trials). **c**, Mean change in DA as a function of port reward probability, separated by rewarded (red) and unrewarded (blue) trials. Changes in DA measured as: peak DA within 0.5s following reward, and minimum DA within 1s following omission, subtracting instantaneous DA at port entry. **d**, Example trace of dLight and running speed across three trials, including when the rat discovered a newly available path (pink star). Scale bars indicate same values as in a. **e**, *Left*, cartoon of rat discovering the absence (*top*) or presence (*bottom*) of a barrier. *Right*, mean DA on each of these trial types; error bands indicate *±* SEM. DA signal is aligned on entry into the hex adjacent to the newly changed hex (pink, newly available; black, newly blocked; each n = 10 rats, 106 events). **f**, Mean change in DA (peak DA within 1s following novel hex discovery – DA 1s before novel hex discovery) was significantly higher for newly available versus newly blocked hexes. Individual rat means are plotted as dots. Bars represent means over rats. * indicates p<0.05, ** indicates p<0.01, *** indicates p <0.001.

We also observed large phasic increases in DA when rats first encountered a newly available hex – i.e., where a barrier had been previously located, but no longer (Fig. 2d-f, p = 1.95*x*10^−3^, two-tailed Wilcoxon Signed Rank test). This was not simply a response to any unexpected sensory event, since encountering a newly *blocked* hex resulted in a consistently smaller or absent DA pulse (2f; available vs blocked: p = * 9.76 10^−4^, one-tailed paired Wilcoxon Signed Rank test; blocked vs 0: p = 0.014, two-tailed Wilcoxon Signed Rank test). Furthermore, the response to newly available hexes was preferentially observed on trials in which rats chose to take the newly open path, rather than ignoring it (Supp. Fig. 2c).

### Dopamine ramps reflect expectations of upcoming reward

We next examined whether the reward-approach ramps previously reported for NAc DA are also present in this more complex spatial environment. Average NAc DA indeed ramped up within each trial, until shortly before arrival at the reward port (Fig. 3a). This overall ramp was significantly positive in nine of ten individual animals (16/19 individual fibers; Supp. Fig. 3). To better understand the computations that give rise to this ramp, subsequent analyses focused on those nine rats. The magnitude of the DA ramp scaled with the current reward probability of the approached port (Fig. 3b), consistent with DA tracking the rats’ evolving expectations of reward. We therefore assessed how DA ramping during port-approach is affected by whether that port was rewarded or not at the last visit (Fig. 3c). DA was generally higher along the whole ramp when the destination port had been most recently rewarded, and lower following an omission. To rule out non-specific effects of recent rewards on DA signals, we performed a multiple regression analysis comparing the impact of the most recent reward outcome at each of the three ports (Fig. 3d). DA ramps selectively reflected expectation of reward at the end of the specific path taken on the current trial, rather than (for example) tracking overall recent reward rate (34), or the rewards available across both potential destination ports together.

**Fig. 3.**
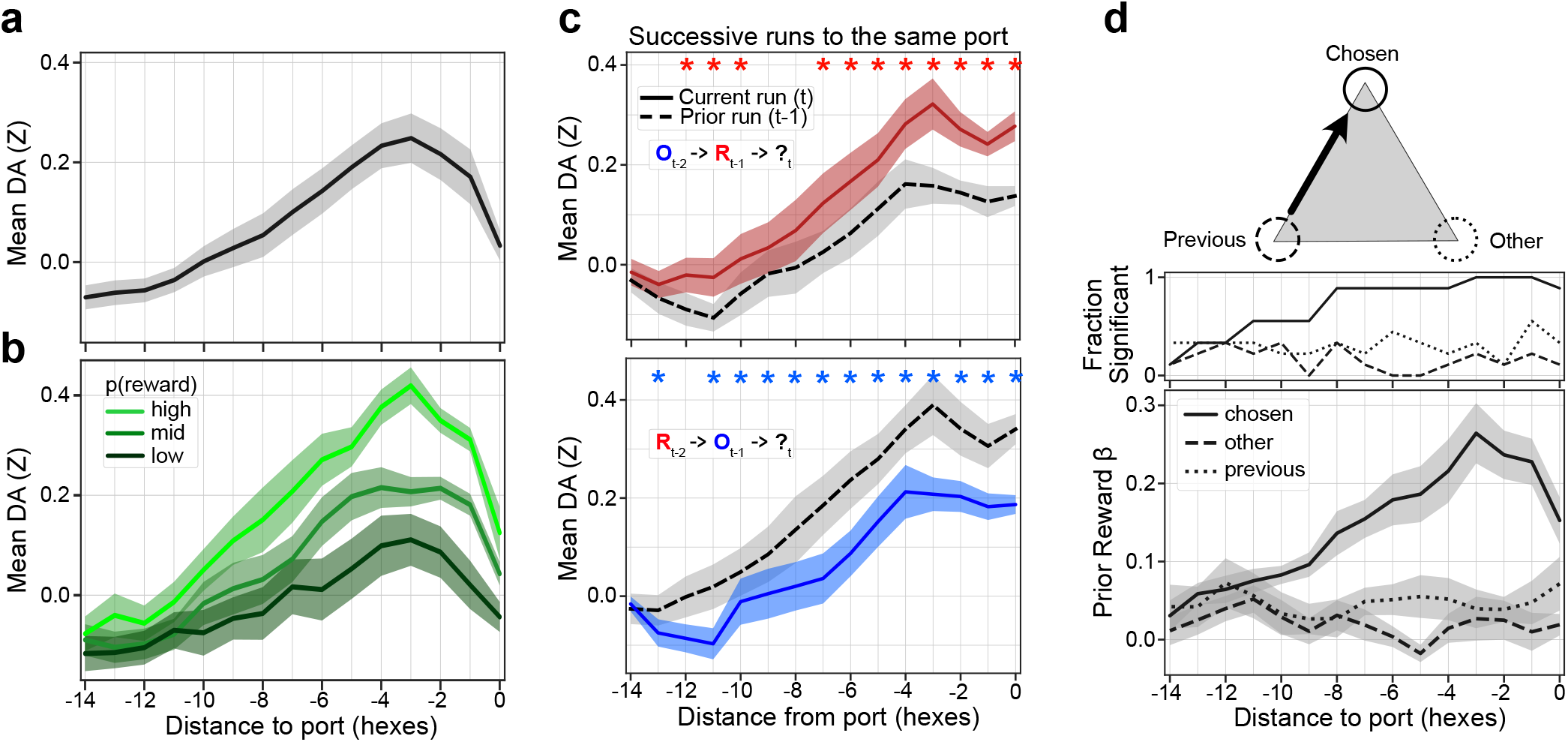
DA ramps reflect dynamic expectations of upcoming reward. **a**, Mean DA as rats approached the reward ports (n = 10 rats, 82 sessions, 15918 trials), as a function of distance. **b**, Mean hex-level DA during port approach, pooled by p(reward) of the destination port. Only the second half (trial number > 25) of each block was included (n = 9 rats, 70 sessions, 7,614 trials). **c**, Examining the effects of reward on DA ramping along successive runs to the same port. Dashed lines indicate the prior run to the port (t-1), and solid lines indicate the current run to the port (t). *Top*, mean DA over successive runs to the same port, where reward was omitted two visits ago (t-2), but reward was delivered the prior visit (t-1; n=9 rats, 1935 sequences). Red asterisks indicate a significant increase in hex-level DA (p < 0.05, one-tailed Wilcoxon signed rank test). “R” and “O” denote rewards and omissions, respectively, on the t-n previous visits to the port. *Bottom*, same as top but examining the effects of a reward omission on the last visit. Blue asterisks indicate significant DA decrease (p < 0.05, one-tailed Wilcoxon signed rank test; n=9 rats, 1909 sequences). **d**, *Top*, maze cartoon illustrating the chosen, other, and previous reward ports for an example trajectory through the maze. *Bottom*, multiple-regression weights for the prior reward outcome at the chosen, other, and previous reward ports as effects on the DA signal (n = 9 rats, 13,448 trials; regressions performed independently for each rat; plot shows mean effect over rats). *Middle*, fraction of rats with significant relationships (non-zero regression coefficient, two-tailed t-test) between prior reward and DA. All error bands show *±* SEM.

### A spatial map of value

These ramping dynamics suggested that DA may signal the evolving, spatially discounted reward expectations throughout the maze environment. We therefore turned to estimating these reward expectations locally: at entry into each hex, from each direction (126 distinct hex-states). As a first pass at estimating these hex values, we again applied a simple learning algorithm to track experienced reward probabilities at each port (Fig. 4a), but then distributed these values, discounted by spatial distance, throughout the maze (“value iteration”; Fig. 4b, (1, 35); see Methods). The resulting hex-level pattern of value closely resembled DA on each trial (Fig. 4c), and a mixed-effects multiple regression analysis revealed a highly significant relationship between DA and these hex values (Fig. 4d; p = 0.000, Likelihood Ratio Test, chi-square distribution with 77 degrees of freedom to account for each session-optimized *γ* value; see Methods). This regression analysis also included running speed, yet hex values accounted for much more of the explained variability in the DA signal (Fig. 4e).

**Fig. 4.**
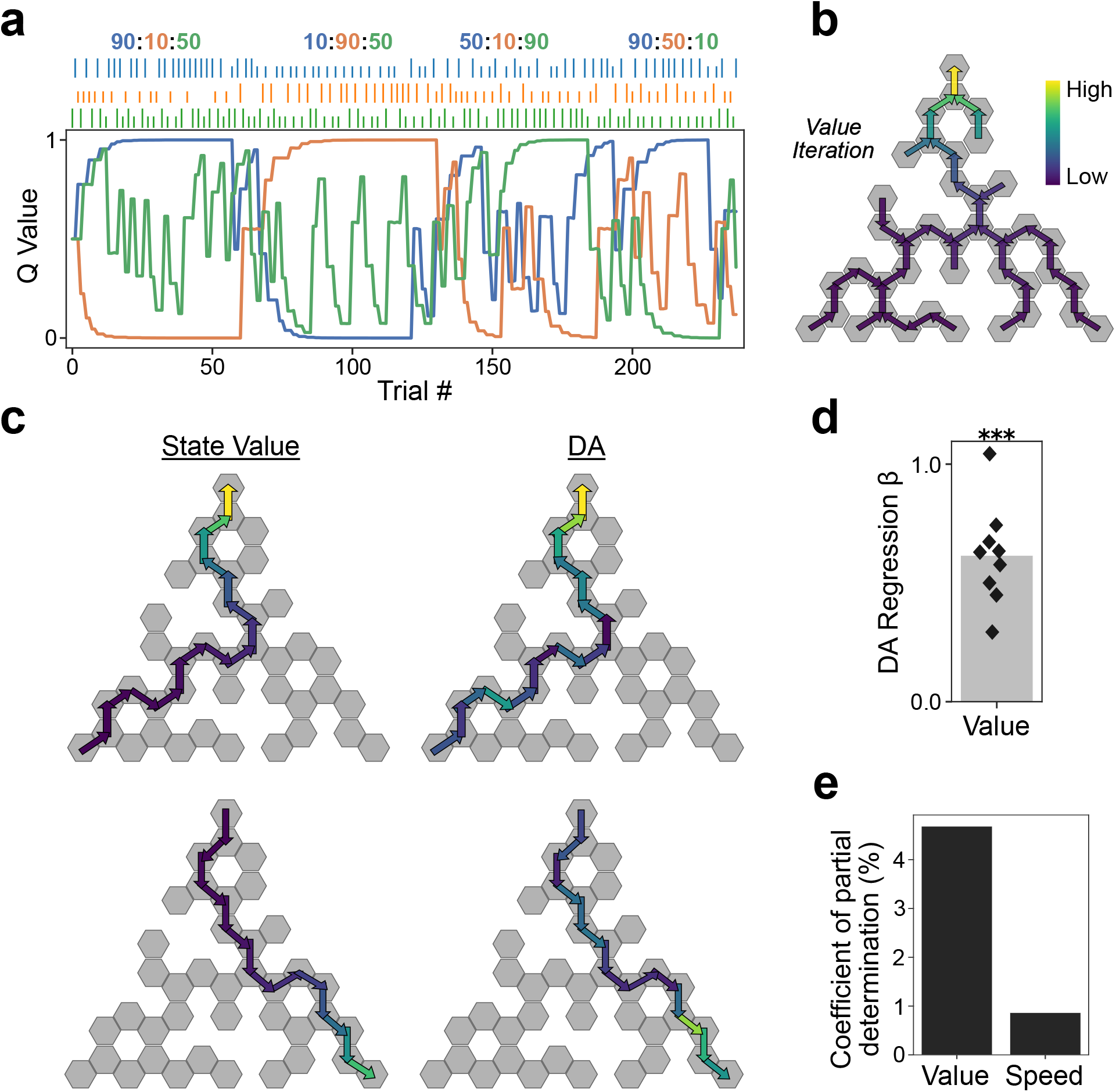
NAc DA reflects a map of place values. **a**, Example of one session showing trial-by-trial evolution of port (Q) values. Numbers at top indicate nominal reward probabilities for the three ports (each in a different color to represent [top : bottom left : bottom right] ports), while tick marks indicate reward outcome on each trial (tall = rewarded, short = omission). **b**, Example value-iteration result from a single trial, spatially discounting the destination port’s Q value over all hexes. Arrows point towards the destination port, and values are defined at entry into a specific hex from a specific direction. **c**, Predicted value (*left*) and observed DA (*right*) during two runs through the maze in one block from the session in panel a. Top example uses the same value map as b. **d**, Regression coefficients for hex value from a mixed-effects regression predicting hex-level DA (n = 9 rats, 77 sessions, 13381 trials, 230,252 hex entries). Bar shows fixed effect over rats; diamonds show fixed effect for each rat over sessions. **e**, Regression model’s coefficients of partial determination for value and running speed. * indicates p<0.05, ** indicates p<0.01, *** indicates p <0.001.

### Over repeated trials, DA signals propagate backwards along taken paths

This value map provides a reasonable first approximation to DA signals as rats run through the Hex Maze. However, the value iteration algorithm requires perfect knowledge of current maze structure, together with the immediate and complete distribution of value updates to all hex-states on every trial. Rat brains might actually use less computationally demanding algorithms to generate place values, and these algorithms could produce tell-tale signatures in value coding while foraging.

In particular we looked for evidence of TD learning, as this has been an especially prominent framework for interpreting DA signals in simpler settings. In its most basic form, TD(0) (also called “one-step” TD), RPEs update only the values associated with the immediately preceding state (1) (Fig. 5a). Therefore, when a sequence of states results in an unexpected reward, earlier states in the sequence do not receive value updates right away. Instead, updates progressively propagate backwards along the sequence, over multiple episodes of experience. A key signature of this type of learning rule is that values of states more distant from reward depend on reward outcomes in the more distant past, rather than the most recent outcomes. TD can also propagate value more rapidly by maintaining memory traces for recently visited locations and using these to determine eligibility for later value updates. Such an algorithm is referred to as TD(λ) (1, 7, 35). By altering the eligibility trace decay parameter, λ, updates can be restricted to the single preceding state (λ = 0, as above), or, at the other extreme, cover the entirety of the experienced path (λ = 1).

**Fig. 5.**
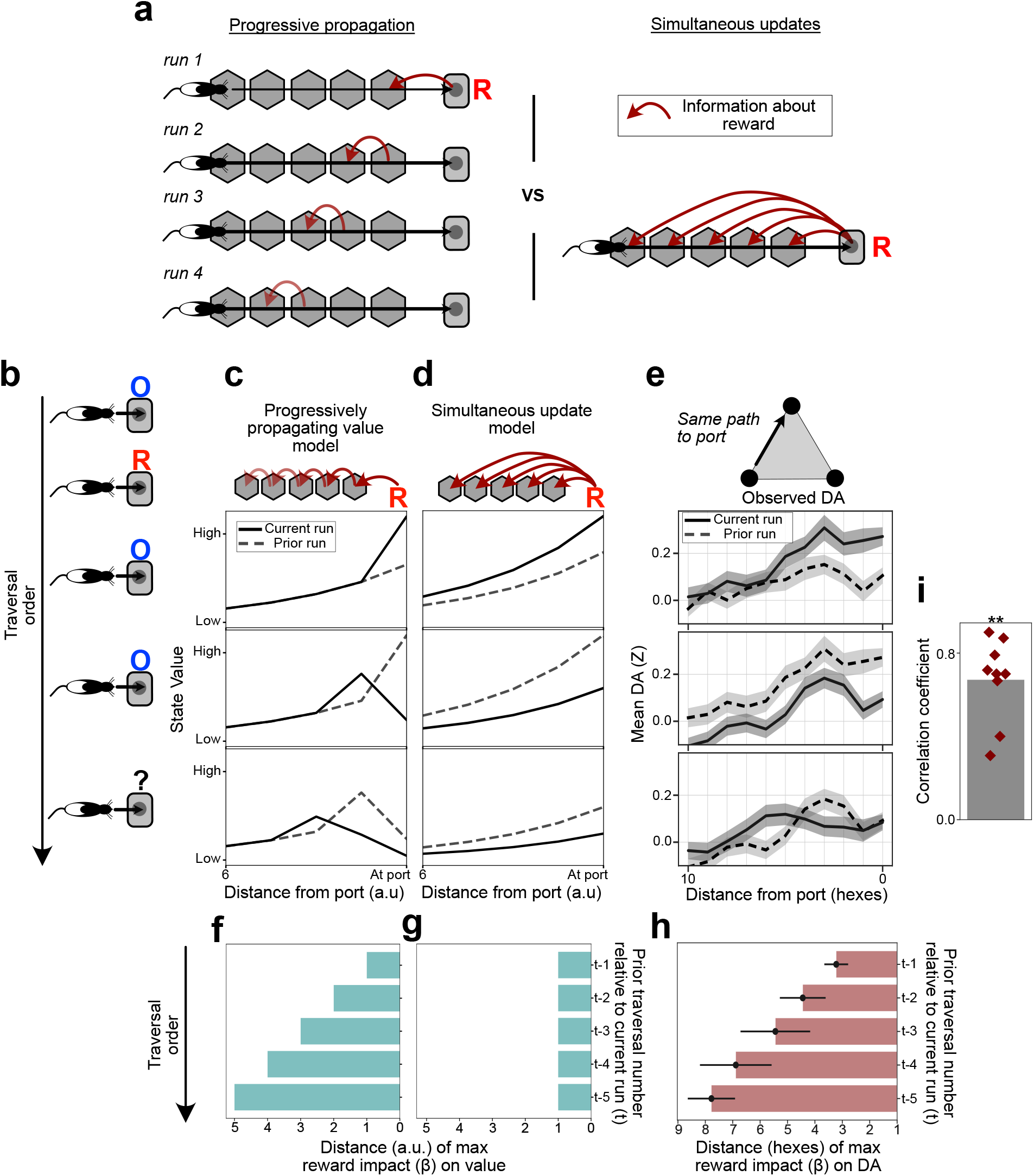
TD-like propagation of DA signals across space. **a**, Cartoon contrasting propagating versus simultaneous value-update algorithms. *Left*, in TD(0) the impact on value coding of a reward on run 1 progressively moves back along the state sequence over successive runs. *Right*, in TD(1) a single reward immediately updates values for all states experienced in the current trial. **b**, Reward sequence illustration over successive runs to the same port, focusing on a single reward among a series of omissions. **c**, Value function from a simulated TD(0) learner over the three last traversals of the sequence in b. Solid lines indicate value function during the current run; dashed lines show value function during the previous run, to illustrate changes. **d**, Value function from a TD(1) learner over the same three sequential traversals. **e**, Observed mean DA traces for the corresponding trial order (i.e. as shown in c; 9 rats, n = 247 trial sets; error bands indicate *±* SEM). **f-h**, Analyzing the distance from the terminal port in which prior rewards have their strongest impacts (linear regression weights) on state value. **f**, Predictions from the TD(0) algorithm over 1000 simulated successive traversals of the same path, with rewards delivered randomly at 50% probability. **g**, Same as f for the simultaneously updating TD(1) learning algorithm. **h**, Results from the same analysis on DA over all successive traversals of the same path (n = 9 rats, 13,427 trials, 235,524 hexes). Bar plot shows mean effect over rats ± SEM. **i**, Correlation between the distance of the peak reward effect on DA (in hexes) and the prior traversal number (1-5 previous traversals). Bar shows mean over rats; diamonds show individual rat coefficients (p = 3.90*x*10^−3^, two-tailed Wilcoxon Signed Rank test).

The resulting difference in value dynamics can be clearly illustrated by considering the impact of a single reward, among a series of omission trials for the same path (Fig. 5b). In simulations (see Methods), with TD(0) the reward evokes a value bump that propagates backwards over the course of multiple traversals (Fig. 5c). By contrast, with TD(1) value is immediately updated across the full traversed path, so that outcomes simply change the gain of the ramping value function (Fig. 5d). We examined DA signals under the corresponding maze conditions: when rats experienced a rewarded trial among a series of omissions for traversals of the same path. The reward appeared to cause a spatial bump in DA, that moved further back from the reward port over successive traversals (Fig. 5e) – i.e., the key signature of TD learning with low λ.

To broaden this analysis to include all sequences of reward outcomes, we turned to multiple regression. We examined how values at each location along a path depend upon reward outcomes on the previous five times this path was taken. As expected, in a TD(0) simulation the information from older reward outcomes had its strongest influence on value farther away from the reward port (Fig. 5f), in stark contrast to TD(1) (Fig. 5g). The same analysis applied to DA signals resulted in a pattern resembling TD(0): older outcomes had the largest influence on DA signals farther from the reward location (Fig. 5h,i; two-tailed Wilcoxon Signed Rank, p = 3.90*x*10^−^3). This provides clear evidence that updates of DA value signals incorporate TD(0)-like progressive, backward propagation.

### DA place values are also globally updated through inference

However, other observations suggested that this TD learning, by itself, provides an incomplete account of value-guided decision-making in the Hex Maze. First, we showed earlier that individual reward outcomes shift DA ramps up or down across a broad path extent (Fig. 3c), not just the hexes close to the port. While both state-to-state chaining and longer-range updates separately arise as the extreme cases of TD(λ), there is no intermediate setting of that model’s parameters at which both of these patterns co-occur. Consistent with this, fitting a TD(λ) hex-state RL algorithm (see Methods) to the observed DA data could model these broad shifts, but the resulting large λ failed to also reproduce the progressive propagation of DA and its dependence on reward history (Supp. Fig. 4a-e). Conversely, removing the broad shifts (by modeling them as a linear scaling of the average DA ramp; Methods) left a residual DA signal that seemed to propagate backwards along the path over trials (Supp. Fig. 4f-h). These results suggest that ramping DA value signals are updated by at least two mechanisms – a TD-like process responsible for backwards signal propagation, and a second process capable of shifting the whole ramp at once.

Second, the behavioral choices of the rats were more sophisticated than would be expected from MF TD alone. In the Hex Maze, each reward port can be reached from multiple starting points (Fig. 6a). MF TD learning would only update values along the path that was actually taken. However, we found that reward at a given port increased the likelihood of choosing that same port at a rat’s next opportunity, *both* when the rat previously took the same path (p = 9.77*x*10^−4^, two-tailed Wilcoxon Signed Rank test) or an alternative path (p = 2.93*x*10^−3^, two-tailed Wilcoxon Signed Rank test) to that port (Fig. 6b). A potential confound could arise from correlations between the most recent reward outcome and prior reward outcomes at the same port, for which the rat may have taken the same path. To control for this, as well as any turn-direction bias, we conducted a mixed-effects multiple regression analysis and included the past five reward outcomes as features (see Methods). We confirmed that a previous reward at a port made current choice of that port more likely, both when the rat had taken the same path to obtain reward (p = 9.47*x*10^−7^) or an alternative (p = 0.0269; Fig. 6b). This suggests the use of model-based (MB) algorithms to infer that hexes along alternative paths to that same reward location have also changed value.

**Fig. 6.**
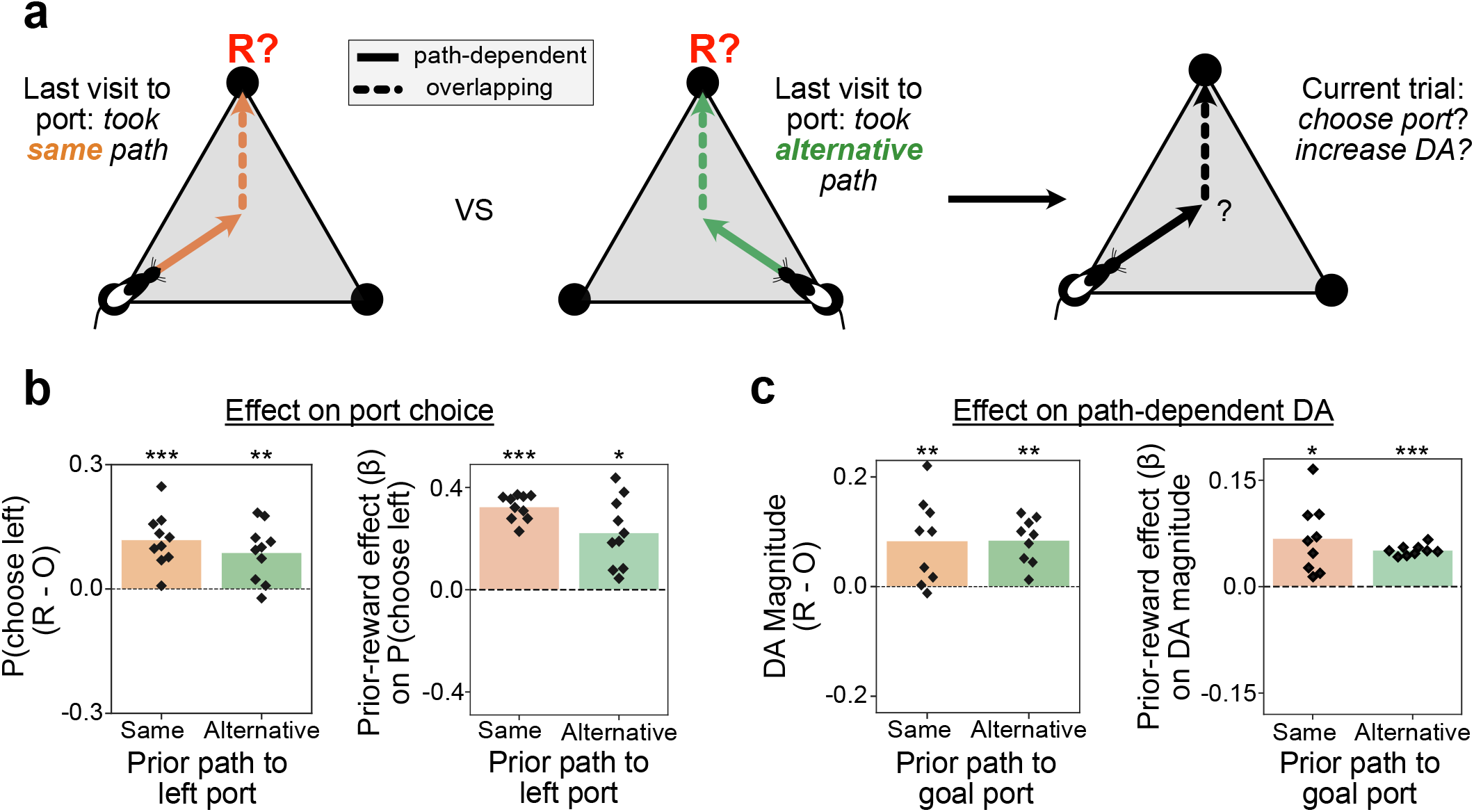
Model-based inference globally updates DA place values and guides choices. **a**, Cartoon of two distinct routes a rat could have taken on the previous visit to a reward port (top of triangle). Portions of each route are distinct based on the starting location (path-dependent hexes; solid line), while other portions overlap (dotted line). **b**, *Left*, the probability of choosing the left (counterclockwise) port after a reward compared to an omission. Separated by trials where, the last time that port was visited, the rat took the same or alternative path. Bars show aggregate means, points show individual rat values (n=10; 2823 rewarded trials along same path, 2439 omission trials along same path, 1799 rewarded trials along alternative path, 1433 omission trials along the alternative path). *Right*, results from two separate regression analyses assessing the probability of choosing the left port as a function of the prior reward at that port. Separated by trials where the rat took either the same (n = 10 rats, 5262 trials) or alternative path (n = 10 rats, 3232 trials) on the last visit to the left port. Regression assumed random effects across rats and estimated a fixed overall estimate for each feature. **d**, *Left*, DA magnitude in path-dependent hexes following a reward compared to an omission, the last time the destination port was visited from either the same (n = 9 rats, 2500 rewarded, 1752 omission trials) or alternative path (n = 9 rats, 1790 rewarded, 1337 omission trials). *Right*, results from two separate regression analyses assessing path-dependent DA as a function of the prior reward at that port, if the rat previously took the same (n = 9 rats, 4252 trials, 33,728 hexes) or alternative path (n = 9 rats, 3124 trials, 32,212 hexes). Regression assumed random effects across rats and estimated a fixed overall estimate for each feature. * indicates p<0.05, ** indicates p<0.01, *** indicates p <0.001.

We therefore assessed whether DA ramps similarly rely upon MB processing and knowledge of maze structure. We confined this analysis to the critical “path-dependent” hexes – those that have no overlap with other paths to the same port (Fig. 6a). We found that prior reward at a port results in elevated DA in these path-dependent hexes, both when the rat previously took the same path (p = 5.86*x*10^−3^, two-tailed Wilcoxon Signed Rank test) or an alternative path (p = 1.95*x*10^−3^, two-tailed Wilcoxon Signed Rank test) to that port (Fig. 6c). Once again, to control for the possibility that this result reflects experiences on even earlier trials, we ran a regression analysis and included the prior five reward outcomes (see Methods). DA still displayed a significant relationship with the most recent reward outcome at the goal port, both when the rat previously took the same path (p = 0.012) or an alternative path (p = 6.49*x*10^−7^) to the goal port. (Fig. 6c). Thus, NAc DA signals reflect MB calculations of inferred future reward from any location, in addition to the MF TD-like learning from direct experience.

### Dual processes account for NAc DA signals during goal approach

To confirm that DA signals are best modeled as arising through the combination of MB and MF learning mechanisms, we applied a dual-process hex-level RL algorithm (Fig. 7). This RL agent experienced the same sequence of hexes and rewards as each rat, and generated corresponding value estimates at each moment. Upon each transition between hexes, MF TD (λ = 0) locally updated just the value of the previous hex-state. The second, MB, process updated the values of all hexes throughout the maze, each time a reward port was visited. This global update reflected the rats’ evolving experience of maze structure, maintained as a recency-weighted average of the tendency of each hex to be followed by a visit to each specific port (whether rewarded or not; Fig. 7a; see Methods). Regression analysis revealed a significant relationship between values in this dual-process model and observed DA (mean β = 0.97 *±* 0.088 SEM, p = 3.73*x*10^−28^, Wald test, z = 11.0).

**Fig. 7.**
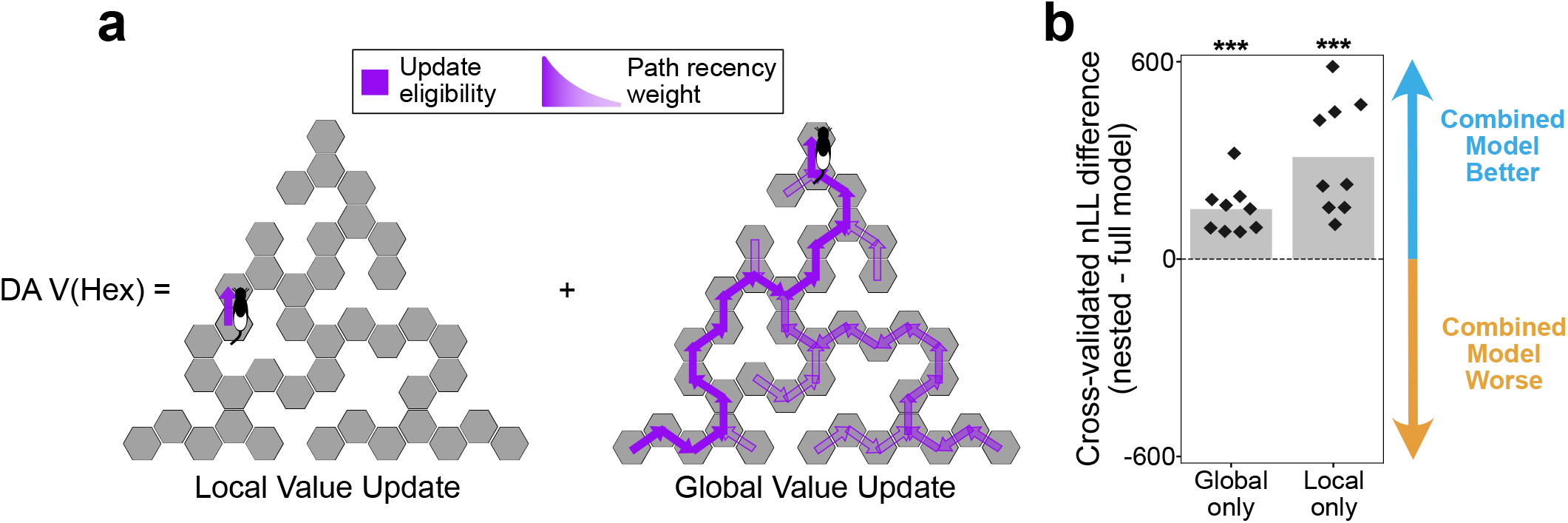
A combined MF/MB model accounts for hex-level DA. **a**, Illustration of the dual-component hex-value RL model’s local and global learning algorithms. *Left*, TD(0) updates the value of the rats’ experienced hex-state at each hex transition. *Right*, upon port entry, a model of the maze’s structure (the hexes that led and could have led to the goal port) is used to globally update hex-value estimates. The model is updated each trial as a weighted sum of previously traversed paths to the chosen port; see Methods for details. **b**, Gain or loss of model fit when each learning component is removed from the dual-component model. Diamonds indicate superior dual-component performance (negative log likelihood, “nLL”), compared to single-component, for individual rats. Bars show mean fit comparison over all rats together. Individual fits also indicated that removing either the TD or global components provided a significantly worse relationship to observed DA signals (global only: p = 1.00*x*10^−14^, two-tailed one-sample t-test, t-statistic = -9.951 *±* 4.008; local only: p = 1.00*x*10^−11^, two-tailed one-sample t-test, t-statistic = -8.642 *±* 2.261).

We then compared the performance of the dual-process model in explaining DA signals to two nested models (36), each with one update process removed (by setting the learning rate to zero). The combined model consistently outperformed either process alone (Fig. 7b). Taken together, this series of analyses provides strong evidence that NAc DA is jointly updated using two classes of learning algorithm: chained updates across sequentially traversed states, and maze-wide updates that leverage MB inference.

## Discussion

Theoretical models of reinforcement learning and decision making have very often employed multi-step navigation through simulated mazes to investigate the performance of distinct algorithms (1). As RL models form the standard framework for interpreting DA signals, it is perhaps surprising that the present study is the first - to our knowledge - to examine real-time DA dynamics in a rich and dynamic spatial environment.

Our observation of a DA pulse at reward receipt, scaling with positive RPE, is consistent with standard DA ideas (although it is noteworthy that such pulses were not observed in a prior study using a simpler T-maze - (19)). By contrast we did not expect to see a similarly-sized DA pulse when rats detected a newly opened path. It is natural to interpret this as some form of error signal, but it is not yet clear what type. DA signals have long been associated with novelty and salient events (37, 38), and some theories have argued that DA can signal a range of prediction errors beyond simply RPE (39, 40). However, a newly blocked path did not evoke a comparably sized DA pulse, suggesting that the relevant feature is not simply an unexpected stimulus, or the need to update models of the environment. It appears that the DA pulse is related to the newly discovered opportunity for action (41), perhaps reflecting the value of discovering possible new paths to reward through exploration (42, 43). Specifying the underlying information processing in greater detail will require further experiments.

A major objective of this study was to investigate the ramps in NAc DA release that occur as unrestrained animals approach rewards (5, 16–18). We and others have previously interpreted this ramping DA as reflecting increasing reward expectation (a.k.a. value). Consistent with this, here we found that ramps track the animals’ recent reward history, for example ramping more strongly when the destination port was rewarded at the last visit. As rats ran through the maze, the moment-by-moment DA levels formed a dynamic map of values: expected rewards discounted by distance from the reward port. This result contributes to an ongoing discussion about whether/how DA signals reflect the costs, as well as the benefits, of potential decisions (44). In the Hex Maze, rats clearly treated distance as a cost, as shown by their reluctance to choose paths leading to more distant reward ports. This cost was incorporated into the DA signal through spatial discounting, producing a net value signal potentially useful for governing motivation from each point. This interpretation fits well with observations that lesions of NAc DA shift motivation in cost/benefit decision-making in maze tasks (45, 46) and that boosting DA can immediately enhance motivation to work (17). An interesting question for future studies is how discounting future rewards over space relates to discounting over time, which has been previously reported for DA signals (47) and may involve distinct time scales in different striatal subregions (48).

Alternative accounts have emerged arguing that DA ramps reflect RPE. This is possible under various assumptions: e.g., that values are rapidly forgotten (20), that there are constraints on the functional form by which value decays in space or time (49), or that animals are uncertain about their current state (50). The present study was not specifically designed to test those ideas. Nonetheless the strong correspondence between DA dynamics and upcoming reward estimates observed here make a value interpretation of ramps the most parsimonious. This is in addition to - rather than instead of - the prediction-error coding noted above. Evidence that DA ramping can be mechanistically distinct from RPE coding comes a prior study in which we compared DA cell firing to release (5). Discrete reward cues evoked RPE-encoding burst firing of identified VTA DA cells, and a parallel increase in NAc DA release. By contrast, NAc DA ramps appeared to occur even without any increase in DA cell firing, suggesting a separate process. A similar comparison in the context of the Hex Maze task could be highly illuminating.

Regardless of whether DA ramping signals value in addition to, or as a side-effect of, error signaling, our results provide new insights into the specific algorithms by which these signals are updated. TD learning has been central to models of DA signals for decades (2), but evidence for the signature progressive backward propagation of value over trials has been sparse and mixed (7–9). Our observation of such propagation here may have been aided by our maze design, in which each hex can correspond to a discrete left/right decision point, and may thus be more likely to be treated by the brain as a distinct “state”. Nonetheless, we could not force the rats to treat hexes as states, and indeed inspection of the propagating DA “bump” suggests that the actual spatial resolution employed by the internal TD algorithm may be in the range of 2-3 hexes (Fig. 5).

Although TD learning is visibly present, we also demonstrated that rats additionally assign credit over long distances in a single step and over paths not directly experienced, suggesting they employ internal models of their environment to guide their DA signals and foraging decisions. We cannot currently say, however, exactly *when* they are using such models. For simplicity we simulated MB value updates as occurring when outcomes are revealed at reward ports. This may be the right time: after running along trajectories through mazes, rats often show sharp-wave ripple (SWR) events, in which dorsal hippocampal place cells can replay recently taken trajectories (51). This replay is especially common after reward receipt (52, 53) and has been proposed to update values along the encoded trajectories (51). Replay can also encode alternative potential paths to reward (11, 54, 55), providing a potential mechanism underlying MB inference of updated values (56). Echoing this perspective, recent research in AI has increasingly emphasized the use of models retrospectively for credit assignment (57, 58).

However, MB value updates might instead, or additionally, be occurring in a prospective manner as rats run towards goals (similar to another family of AI algorithms that that use models for planning; (59)). First, SWRs occur not only following reward receipt, but also during pauses in behavior, when place cells can encode locations predictive of the animal’s future path (60). Such “forward” replay toward goal locations could, in principle, accomplish MB value updates similar to “backward” replay from them (56). Additionally, actively running rats show “theta sequences”, in which the maze location encoded by hippocampal places cells sweeps forward ahead of the animal within each theta cycle (61) and can even rapidly switch between representations of distinct possible future paths (62, 63). It may be that theta sequences help retrieve current values of potential goal locations to help guide decision making, although this is not yet known. We note that such cognitively demanding planning processes may be only activated when the brain perceives a need to do so. If DA ramping is linked to prospective planning, this could explain why DA ramps peak just ahead of actually reaching the goal (Fig. 3). Within the last 3 hexes of the path the reward port is directly visible, so no internal calculations are required for navigation. This account is also consistent with prior reports that ramps disappear entirely as rat behavior becomes increasingly routine (22), and can reappear immediately if task contingencies change (18). There is analogous evidence that nonlocal activity in hippocampus declines over repeated experience, including both SWRs (53) and cycling between multiple paths during theta sequences (64). Furthermore, there is substantial behavioral and pharmacological evidence that NAc DA is specifically required when animals need to flexibly calculate trajectories to reward (e.g. from a variable start location) rather than performing a stereotyped sequence of actions (65, 66). For future reports, we aim to combine measurements of DA ramps with high-density hippocampal recordings, to gain greater access to the internal calculations driving DA dynamics during active foraging.

## ACKNOWLEDGEMENTS

We thank Simon Little, Colin Hoy, Vijay Namboodiri, Anna Grzymala-Busse, and members of the Berke Lab for providing valuable feedback on manuscript drafts, Ali Mohebi for initial assistance with fiber photometry procedures, hardware configuration, and surgical procedures, Yang-Sun Hwang for assistance with rat raining and fiber photometry, Lilian Pelattini for assistance histology, and other members of the Berke Lab for assistance and advice. This work was supported by the National Institute on Drug Abuse, the National Institute of Neurological Disorders and Stroke, the National Institute of Mental Health, the State of California, the National Science Foundation, and the University of California, San Francisco (R01DA045783, RF1NS116626, R01NS123516, IIS-1822571).

## AUTHOR CONTRIBUTIONS

T.A.K. developed, built, and optimized the Hex Maze, performed the photometry experiments, analyzed the photometry data, and performed the computational modeling in partnership with N.D.D.. A.E.C. and L.M.F. provided regular scientific input and feedback. N.D.D. provided guidance and code for computational modeling, and developed the computational models with T.A.K.. J.D.B. designed and supervised the study, developed the Hex Maze, and wrote the manuscript together with T.A.K..

## Methods

### Animals

All animal procedures were approved by University of California San Francisco Institutional Committees on Use and Care of Animals. Male (300–650 g) and female (250-400g) wild-type Long-Evans rats were maintained on a reverse 12:12 light:dark cycle and tested during the dark phase. Rats were mildly water deprived, receiving 30 minutes of free water access daily in addition to fluid rewards earned during task performance. No sample size precalculation was performed.

### Behavioral Task

The Hex Maze consists of a 1.30m-per-side equilateral triangular platform with liquid reward ports at each vertex. Solenoid valves control delivery of sucrose solution (10% sucrose, 0.1% NaCl) in 15μL droplets. Infrared photobeam sensors detect entry into the reward ports. To prevent uncertainty over reward delivery, a brief (70ms) 3.0KHz tone was played through a speaker below the center of the maze immediately before solenoid valve opening. Equally spaced columnar barriers divide the maze into 49 hexagonal units (“hexes”). Additional barriers can be placed in any combination of the 49 hexes to create unique maze configurations. The apparatus was controlled by an Arduino Mega, while the Open Ephys software, Bonsai, was used for behavioral and video data acquisition.

Prior to implantation, rats were mildly water deprived and trained in the maze for approximately three weeks. Pre-training consisted of learning to poke into reward ports to receive reward, at 100% delivery probability with no additional barriers. Rats were pretrained until they completed an average of at least one trial per minute in a 60-minute session (1-2 sessions to reach criterion on average). To discourage a sit-and-wait strategy, after each visit to a port that port was not rewarded again until another port is visited (this rule is present throughout training and testing). Rats were then trained on the task until reaching criterion (>= one trial per minute in a 90 to 120-minute session). Before each session, barriers (8 or 9) are added to the maze to create a configuration that is novel to the rat. To prevent clearly visible paths between ports, we ensured that at least one barrier obstructed each direct path. We also configured at least one path to be longer or shorter than another path, to create distinct distance costs associated with different paths. In the probability-change variant, the maze configuration stays consistent throughout a session, but reward probabilities are changed following each block (50-70 trials). Probabilities are reassigned pseudo-randomly, according to the rule that the most rewarding port and the least rewarding port are not the same for two consecutive blocks. In the barrier-change variant of the task, reward probabilities remain fixed throughout the session, while one barrier is moved at each transition between blocks. Upon a block change, barriers are moved strategically to simultaneously alter the lengths of multiple paths: a path that was relatively longer in one block will become relatively shorter in the next, and a path that was relatively shorter in one block will become relatively longer in the next. Barriers were physically moved by the experimenter, who entered the task area after the rat poked into a reward port on the last trial of a block. To prevent the development of associations between experimenter entry and configuration changes, the experimenter randomly entered the task area to briefly raise and lower a barrier – without changing the maze configuration – at least once during each training session. Each daily test session used either the probability-change or barrier-change variant, and we only included behavioral sessions with 100 or more trials for further analysis. Individual rats were also excluded from analysis altogether if logistic multiple regression revealed a non-significant effect of either reward probability or distance cost on their port choices (n=1 rat without significant reward effect, and n=1 rat without significant distance effect from a total initial dataset of n = 12 rats). All rats experienced sessions with port-reward probabilities drawn from a set of [0.9, 0.5, 0.1], but four rats also had probabilities drawn from [0,8, 0.5, 0.2] on a subset of sessions.

Rats’ implant caps were labeled and tracked using Deeplabcut (67). Custom code was used to segment the maze into hexes and classify hex occupancy. For time points with missing position information (i.e., when rat’s heads were momentarily obstructed by barriers), we used the maze’s hex adjacency matrices to interpolate between hexes.

### Fiber photometry

The nucleus accumbens was bilaterally targeted using the following coordinates in relation to bregma: +/-1.7mm medial, 1.7mm anterior, and 6.2mm below brain surface. Virus – 1μL of AAVDJ-CAG-dLight1.3b (Vigene) at a titer of 2×1012 – was delivered using a stereotaxic injection pump (Nanoject III). Virus was injected 200μm ventral to the target coordinates, as described in (5). During the same surgery, 200μm optical cannulae were subsequently implanted and cemented in place. A subset of rats (n = 4; IM-1322, IM-1398, IM-1434, IM-1478), were also implanted with a custom electrophysiology probe in the dorsal hippocampus.

Rats were removed from water deprivation at least 24 hours prior to surgery. One week after surgery, rats began mild water deprivation and were retrained on the task, while waiting for expression of dLight. Rats began photometry recordings in the maze at least two full weeks following surgery. Only one implanted fiber was recorded in a given photometry session.

Photometry data acquisition methods have been described previously (5). Baseline correction was performed using the adaptive iteratively reweighed Penalized Least Squares (airPLS) algorithm (68). Baseline-subtracted 470nm and 405nm (isosbestic control) signals were then each standardized (z-scored) using a session-wide median and standard deviation. The standardized reference signal was fitted to the 470nm using non-negative robust linear regression, and the normalized fluorescence signal was computed by subtracting the fitted reference signal from the standardized dLight signal. To reduce the frequency and severity of optical artifacts, we used a pigtailed optical commutator (Doric Lenses), oriented horizontally, and manually controlled its movement using a custom stepper-motor interface. Recording locations were histologically verified using immunohistochemistry (5). Recording sessions were excluded if a recording failure occurred at any point during the session, such as an optical fiber becoming broken or unplugged.

For all time-based analyses, the dLight signal was downsampled to 250 Hz and smoothed with a rectangular 100ms rolling mean. For hex-level photometry analyses, we calculated the mean dopamine within each traversed hex on a given run. For comparison with RL model variables, we computed mean dopamine within each traversed hex from each possible direction of entry. This included repeat entries into hexes traversed multiple times within a trial (e.g., after leaving a hex, entering a dead end, and running back to through that same hex). To avoid analyzing subsets of data where rats mistakenly returned to the previous (now unavailable) port, only data between the final poke at one port and the first poke in at the subsequent port was included. For event-aligned plots, traces were first averaged over sessions within each rat before taking the average over each rat, unless otherwise specified. Unless otherwise specified, we treated individual rats as the unit of analysis, rather than e.g. fiber recording locations.

### Reinforcement Learning Models

#### Q(port) learning

To estimate the rats’ expected value at each port on each trial, we used a simple, trial-based Q learning algorithm. The model learns values associated with each port using the following update rule:

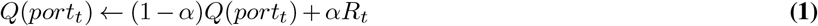

where α is the learning rate, t denotes the current trial, and R denotes reward received at the end of the trial. Choice was modeled as a probabilistic decision between the two available destination ports, left (“L”) and right (“R”), denoted by their position clockwise or counterclockwise from the animal, on each trial using a softmax distribution:

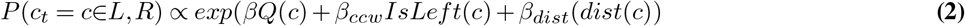

The inverse temperature parameter, β, controlled the degree to which the value of the destination port, Q(port) influenced port choice. The “*β*_*ccw*_” term was added to control for leftward (counterclockwise) turn biases, and a distance-sensitivity (“*β*_*dist*_”) term was added to control for effort cost scaling with the distance dist(c) to the port. “IsLeft” encodes whether the choice, “c”, was leftward from the current port. Parameters were optimized to maximize fit to rats’ observed port choices.

#### Value iteration

We sought to generate spatially discounted chosen value estimates for each hex at the individual-trial level, in a manner faithful to the maze configuration on each trial. We first specified ground truth hex-state transition matrices for each unique maze configuration. We then used a value-iteration (1, 35) algorithm to dynamically estimate state value over each hex-state. Here, hex-states were defined by hex ID (1-49) paired with the direction of hex entry, which resulted in a 126-hex-state state space (each hex has between one and three possible directions of entry). For each trial, the reward function was set to zero at all states other than the chosen port, which was set to the goal port’s Q value on that trial. Hex values were initialized at zero, and value was iteratively learned by taking the maximum of the available discounted next-state values, over all hexes, until convergence. The update rule took the following form:

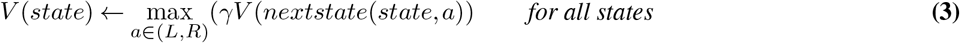

where “a” is a left or right action exit from the current hex-state, and nextstate(state,a) hex-state’ is determined by taking action “a” one step through the transition matrixis the state obtained (through the transition matrix) by exiting state with action a. The discount factor, γ, was optimized for each behavioral session to maximize the fit to DA (minimizing negative log likelihood of the observed DA, given the estimated value function (36)).

#### TD(λ) toy-path value learner

To test distinct predictions about reward propagation over space, we created a simple TD model with an adjustable eligibility-trace parameter (TD(λ) with replacing traces; (Sutton and Barto 2018). Each traversed state was associated with an update eligibility that decayed exponentially – by a factor of λ – with each timestep. To model locally chained value propagation, we implemented a one-step TD model by setting λ equal to zero (TD(0)). To model updating over the entire traversed path, we set λ equal to one. Due to the absence of RPE over successive traversals of the same path under TD(1), value updates only occur at the terminal state, and for the entirety traversed path. Under these conditions, TD(1) is equivalent to a Monte-Carlo learning process (1). Eligibility traces e were initialized at zero, and the update rules were as follows, at each step t:

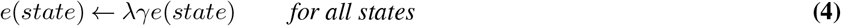

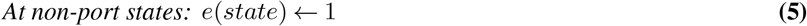

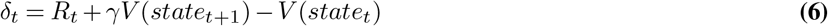

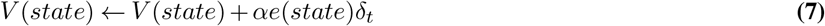

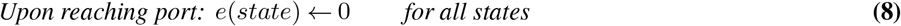

where V is the value function, γ is the discount factor, and α is the learning rate. For clear visualization of model predictions, α was set to 0.8 and γ was set to 0.8. To recreate a ramp similar to the DA signal, each learner started with a baseline value function peaking at 0.4 and discounted by a factor of 0.8. The toy environment was implemented as a six-state sequential path to a reward port, and the reward function equaled zero at all states except the terminal port. Port rewards could be set by the experimenter in order to visualize value functions over specific reward outcome sequences. Alternatively, rewards could be drawn from a random distribution, also defined by the experimenter. For the regression analysis in Figure 5, assessing the relationship between prior reward outcomes and model value estimates at each state, we simulated 1000 trials with random rewards delivered at 50% probability.

#### Dual-component hex-value learner

To compare contributions of spatially local TD and maze-wide inference-based learning processes, we developed a value learning algorithm over hexstates (location and direction, defined as before), with two separate value-update components: local TD(0) value learning, and a maze-wide model-based update.

A one-step TD(0) update occurred at every hex entry according to the following update rule:

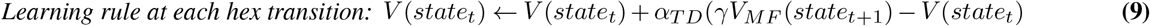

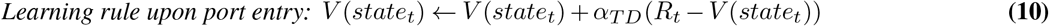

where *α*_*T D*_ is the TD learning rate, and *γV*_*MF*_ is the spatial discount factor. The reward function, R, was zero for all non-port hexes. Hex-state values were initialized at 0.2, to convey a small uniform expectation of future reward from all locations. Upon reaching a reward port, model-based updates were also performed over the entire map upon reaching a reward port according to the following rule:

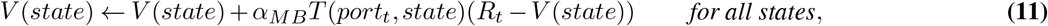

where *α*_*MB*_ is the model-based update learning rate, and T(port,state) weights the update by the discounted on-policy distance from each state to the current port. This map the tendency matrix, T, is a recency weighted representation of all paths that have previously led to the goal port. The tendency matrix is learned online by recency-weighted averaging over states encountered on paths into the port. In particular upon each port arrival, it is updated according to:

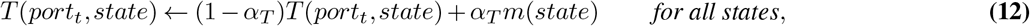

using learning rate *α*_*T*_ and a where the memory trace vector, m, of the most recent path into the port, reflecting each hex traversed on the current trial, discounted by the experienced distance from the port.is a vector of the hexes traversed on the current trial, and *α*_*T*_ is a learning rate. The update weight of each hex is further discounted as a function of timesteps within an experienced path to the goal port. The memory trace m is itself initialized to zeros at the start of each trial, then learned over the trial by discounting and accumulation at each timestep. It learns this representation of distance from the reward port through the following discount and accumulation rules at each timestep:

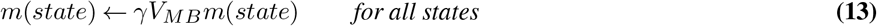

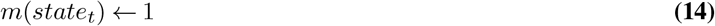

In this way, T reflects a model-based expected eligibility trace for possible paths to the port, comprising both experiential eligibility from the just-completed path into the port (analogous to TD(1)), and counterfactual eligibility arising from a recency-weighted average over previous port entries (57, 69).

#### Hex-state value TD(λ) learner

We also considered an alternative model for learning hex-state values, based on TD(λ). This algorithm maintained an eligibility trace of recently visited hex-states to propagate updates backwards at each timestep. By optimizing the trace decay parameter, λ, to fit the observed DA at each timestep, we could estimate the spatial extent of value updates, on average. Value learning was implemented according to the following rules:

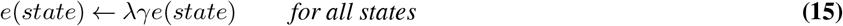

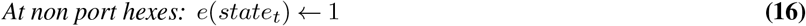

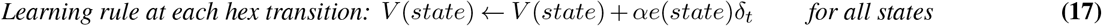

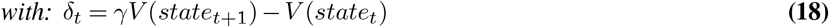

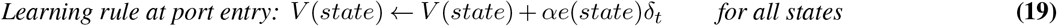

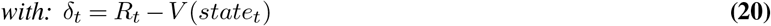

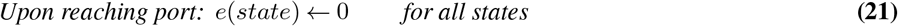

#### Dopamine regression

We combined each learning model with a linear regression observation function to model the dopamine timeseries, i.e. *DA*_*t*_ = *β*_0_ + *β*_*V*_ *V* (*state*_*t*_) + *ϵ*_*t*_, with noise *ϵ*_*t*_ ∼ *N ormal*(0, *σ*^2^). Here, the parameter *β*_*V*_ captures any covariation between modeled value and the measured dopamine timeseries.

#### Model fitting

We optimized the free parameters of the learning algorithms by embedding each of them within a hierarchical model to allow parameters to vary from session-to-session. Session-level parameters were themselves modeled as arising from a population-level Gaussian distribution over sessions, across rats. We estimated the model, to obtain best fitting session- and population-level parameters to minimize the negative log likelihood of the data using an expectation-maximization algorithm with a Laplace approximation to the session-level marginal likelihoods in the M step (70). For hypothesis testing on population-level parameters (*β*_*V*_), we computed an estimate of the information matrix over the population-level parameters, taking account of the so-called “missing information” due to optimization in the E-step (71), itself approximated using the Hessian of a single Newton-Raphson step.

For the value-iteration algorithm, which only sought to estimate the discount factor, γ, we used a simpler function-minimization protocol. On a session-by-session basis, we found the minimum of the negative log likelihood function of the DA data, given γ. As this was a simple scalar function, we used the minimize_scalar function from the SciPy package in Python. Parameter search was unbounded using Brent’s algorithm, but γ values were rescaled between 0 and 1.

#### Model comparison

To isolate the contributions of each independent learning component, we created two nested models: one with *α*_*T D*_ and *γ*_*MF*_ both set to 0 (MB update only), and another with *α*_*M*_ *B, α*_*T*_, and *γ*_*MB*_ all set to 0 (TD update only), and we compared each of these to the full model. In order to compare models with different numbers of free parameters, correcting for any bias due to overfitting, we computed a cross-validated approximation to the negative log marginal likelihood for each session (36). Specifically, we used leave-one-session-out cross validation for the population-level prior parameters and a Laplace approximation for the per-session parameters: for each session, we refit the population-level model omitting that session, then conditional on that prior, we computed a Laplace approximation to that session’s log marginal likelihood. We aggregated these per-session scores to obtain a total score for each rat and model. Finally, we use paired tests on these scores across rats, between models, to formally test whether any model fit consistently better over the population of rats. We depict relative fit subtracting out the dual-component model fit scores, so that positive values indicate superior dual-component model fit.

## Data Analyses

### Port-choice analyses

The frequencies of port visits and path choices were calculated using a five-trial rolling mean. To compute changes in visit frequency, we subtracted the mean frequency from the five trials prior to a block change from the frequencies after a block change. Note that paths here, and in most analyses, are defined by port visits (e.g., running from port A to port B), rather than specific sequences of hexes. “Better” and “Worse” ports were defined as those where the reward probability increased or decreased, respectively, compared to the prior block. This included changes from 10% reward probability to 50% reward probability, so the “Better” port was not necessarily the highest reward probability port in the maze. Similarly, “Longer” and “Shorter” paths were defined relative to the previous block, and paths whose length did not change were not included in this analysis.

All mixed-effects regression analyses were performed in R using the package lme4. Random effects were estimated over the levels of rat and session-within-rat. To identify any significant contributions of reward probability and path length on choice, we used a logistic mixed-effects regression of the following form:

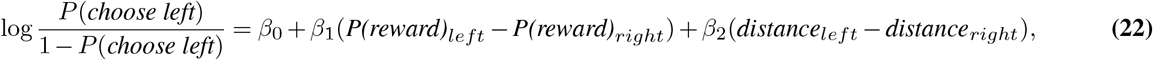

where the intercept captured any variation due to turn-direction bias. “Left” was defined on a trial-by-trial basis as the left of the two available ports, when oriented away from the previously visited port. For example, if the top port had just been visited, the bottom right port would be left, and the bottom left would be right. To avoid periods when rats are learning the probabilities of reward, we only included data from the second halves (trial number > 25) of each block. Both probability differences and length differences were scaled between zero and one to compare effects in common units.

To isolate any effects of inference on port choice, we ran a similar logistic mixed-effects regression of port choice:

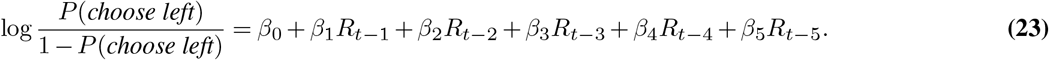

where “t-n” denotes prior trials where the left port was visited, and R denotes the reward outcome on that trial. Critically, we ran this regression for two subsets of data: trials where the rat took the same path to the goal port the last time it was visited, and trials where the rat last took an alternative path to the goal port. Paths, here, were defined based on the start and end ports, not the specific sequence of individual hexes traversed.

In addition, we sought to avoid possible confounds that arise due to decaying reward representations over time. For example, for a port that has not been visited in 10 trials, memory of the last outcome may have decayed, or uncertainty may have increased, compared to a port visited one trial ago (i.e., when a rat has been running back and forth between two ports and ignoring the other). To control for variations in the trial-lag length between traversals to the port of interest (the left option), we only included trials where the left available port was visited exactly two trials prior. This way, we are not comparing results from recent same-path reward to older alternative-path rewards, or vice versa.

### Ramp analyses

Ramp slopes were estimated by fitting a linear regression model to the hex-level DA along the last 15 hexes traversed before port entry, in each session. To test for significance, average ramp slopes were first computed for each session, and then a two-tailed Wilcoxon Signed Rank test assessed whether a rat’s session-average slopes were significantly different from zero. If a rat did not show significant positive ramping (n=1) they were not included in remaining analyses of DA ramping and value coding.

To scale and remove average ramps from individual-trial DA traces, we first calculated the average ramp over the last 10 hexes traversed for each rat. Because we were interested in scaling the entire ramp as a function of estimated gain, we needed to remove any negative values. To do this, we first rescaled each rat’s average ramp between 0.1 and 0.9 (we refer to this as the control ramp, for clarity). For each path traversal of interest, we then fit a linear regression of the observed DA data to that rat’s control ramp. An intercept captured remaining broad directional differences in the ramp (e.g., when the initial portion of the observed DA ramp was negative). We then scaled the control ramp by the estimated regression coefficient, added the intercept to the scaled control ramp, and subtracted this result from the DA trace. We were left with residual DA values, which we used for visualization in Supp. Fig. 4g-h.

### Barrier-change dopamine analyses

To analyze discovery of a barrier change (either newly available or newly blocked) we aligned signals on first-detected entry into a hex immediately adjacent to the changed hex. At these hex transitions, the changed hex is readily visible. Initial new-hex exposures where the rat subsequently entered the new path were defined as those where the rat entered the newly available hex directly following its discovery.

### DA regression analyses

We needed to isolate the hexes where values will differ depending on experience-based versus inference-based updates. To this end, we excluded all overlapping hexes between the same and alternative paths to the goal port. In other words, we only included the hexes prior to the first choice point on each trial where the rat has the opportunity to choose between the two available ports (see Fig. 6).

To assess whether DA reflected the last reward outcome at the goal port following a traversal of the same path-dependent hexes and/or an alternative sequence of hexes, we ran a mixed-effects regression of the following form:

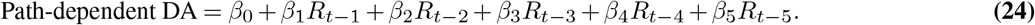

where “t-n” denotes prior trials where the goal port was visited, and R denotes the reward outcome on that trial. Similar to the port-choice analysis, we ran this regression for two subsets of data: trials where the rat previously took the same path to the goal port, and trials where the rat took an alternative path to the goal port. Again, to control for biases that can arise due to differences in the number of trials since the port was last visited, we exclusively analyzed trials where the goal port was visited two trials ago. The inclusion of the prior five outcomes at the goal port controlled for DA scaling effects due to earlier rewards at the same port.

**Supplementary Figure 1.**
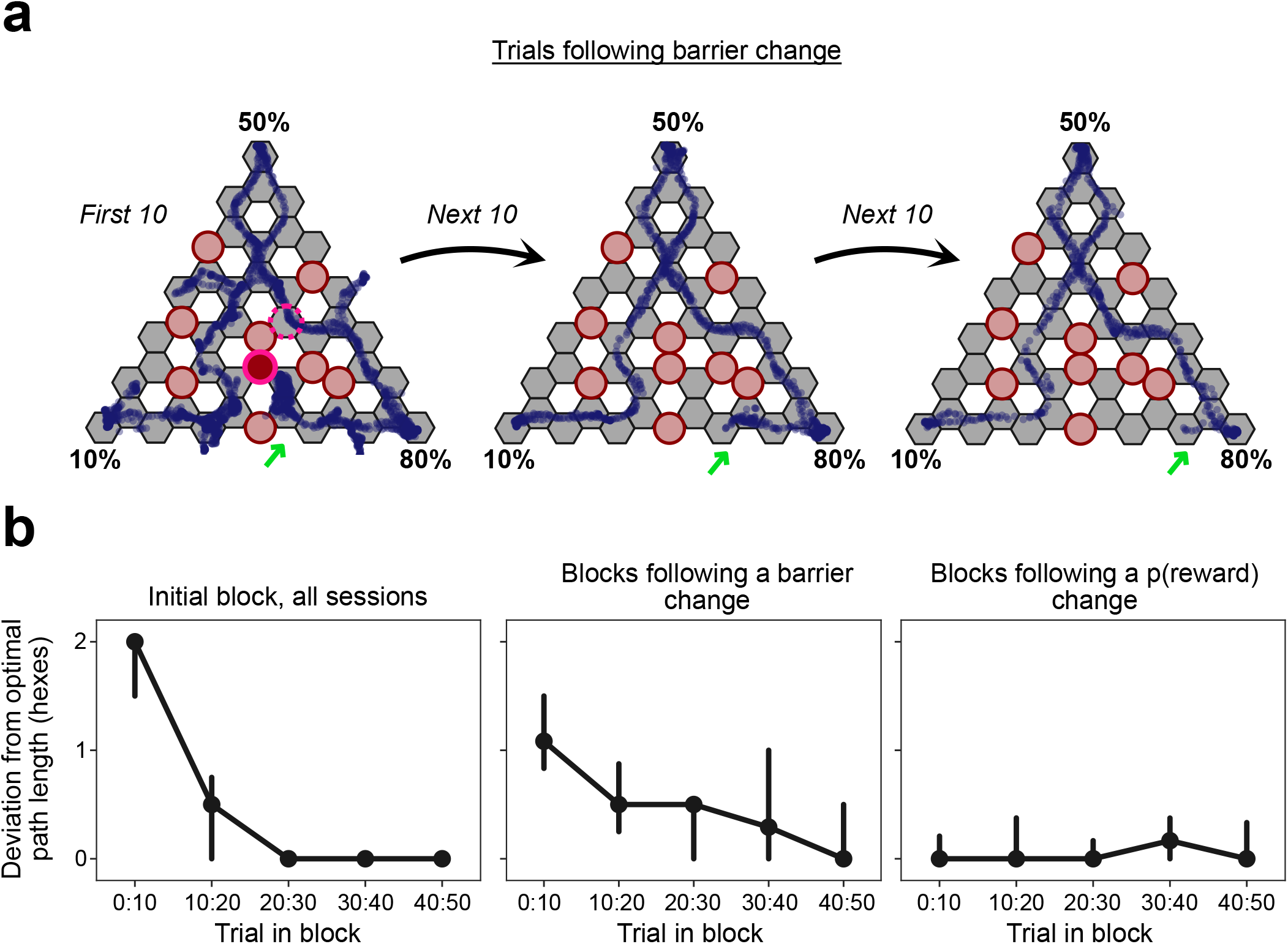
Navigational adaptations to Hex Maze configuration changes. **a**, Position data from the first, second, and third group of ten trials following a barrier change. Green arrows highlight the progressive reduction in the distance traveled into a novel dead-end path. Barrier change is from the same block as Fig. 1d. **b**, Deviation from the optimal (shortest) path lengths over the course of the initial blocks of all sessions (*left*), blocks following a barrier change (*middle*), and blocks following a p(reward) change (*right*). Deviation is measured as the number of extra hexes traversed beyond the shortest possible path length. Dots show median values in ten-trial bins, error bars show 95% confidence intervals.

**Supplementary Figure 2.**
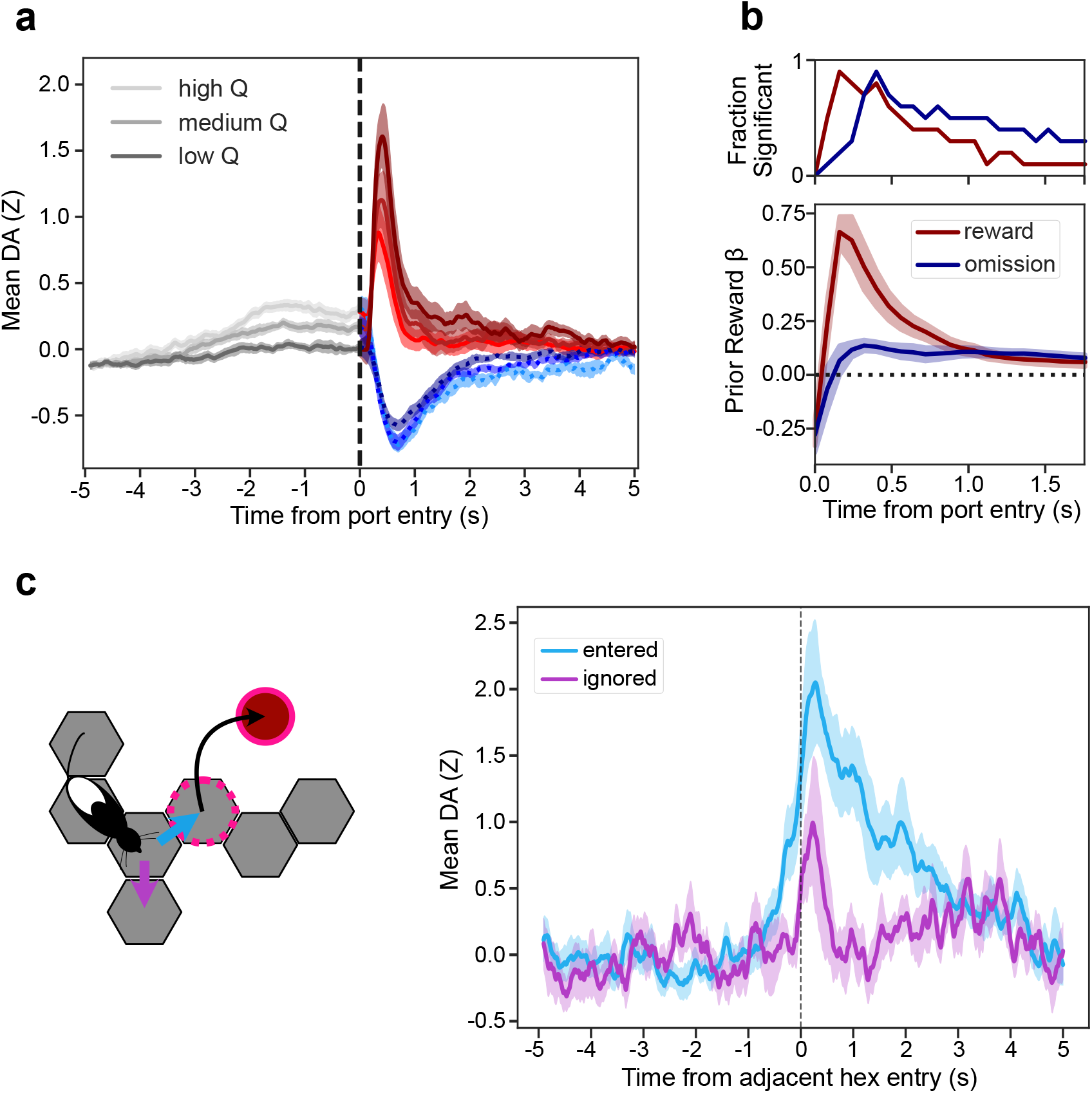
Extended analysis of DA pulses. **a**, Port-entry aligned DA in the second half (trial number > 25) of each block (n = 10 rats, 82 sessions, 9,079 trials), pooled by terciles of the RL model Q value for the chosen port. **b**, *Bottom*, regression weights for Q-value-derived RPE (see Methods) on DA following port entry (100ms bins over the first 2s). Separate regression weights are shown for RPE following reward (red) and omission (blue). Regressions were performed independently for each rat. *Top*, fraction of rats with a significant relationship (non-zero regression coefficient, two-tailed t-test) between RPE and DA in the time bin. **c**, *Left*, cartoon of rat choosing to enter (light blue) or ignore (violet) the newly available path. *Right*, DA aligned on entry into the hex adjacent to a newly available hex, broken down by whether the rat subsequently entered (n = 9 rats, 77 events) or ignored (n = 7 rats, 25 events) the novel path. Four rats either never chose to enter the ignored path option, or entered on fewer than three instances in total, so they were excluded from analysis. All error bands indicate *±* SEM.

**Supplementary Figure 3.**
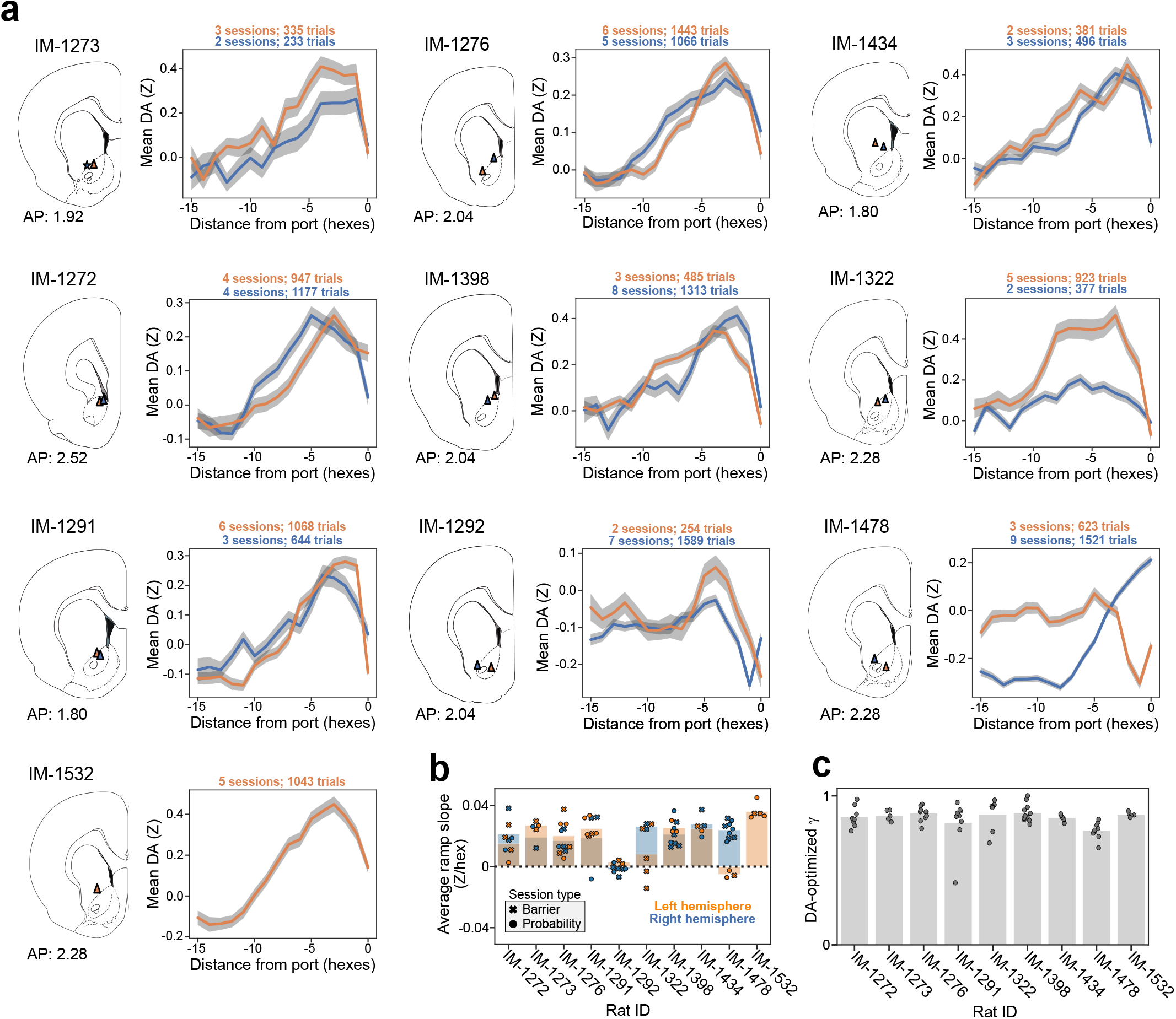
Individual-animal recording locations and DA goal-approach ramps. **a**, Histologically identified fiber locations for each animal, paired with the average ramp *±* SEM for the corresponding implanted fiber. Orange denotes right hemisphere and blue left. Blue asterisk signifies the inferred fiber location of an implant with missing histology (IM-1273 left hemisphere). **b**, Average ramp slope for each recording location (bar plot), with individual sessions marked as “x” for left and “o” for right hemisphere. **c**, Discount factors (γ) for each recording location with a significant ramp, fit to the observed DA data in the value-iteration algorithm. Bars show rat means, and dots show session values.

**Supplementary Figure 4.**
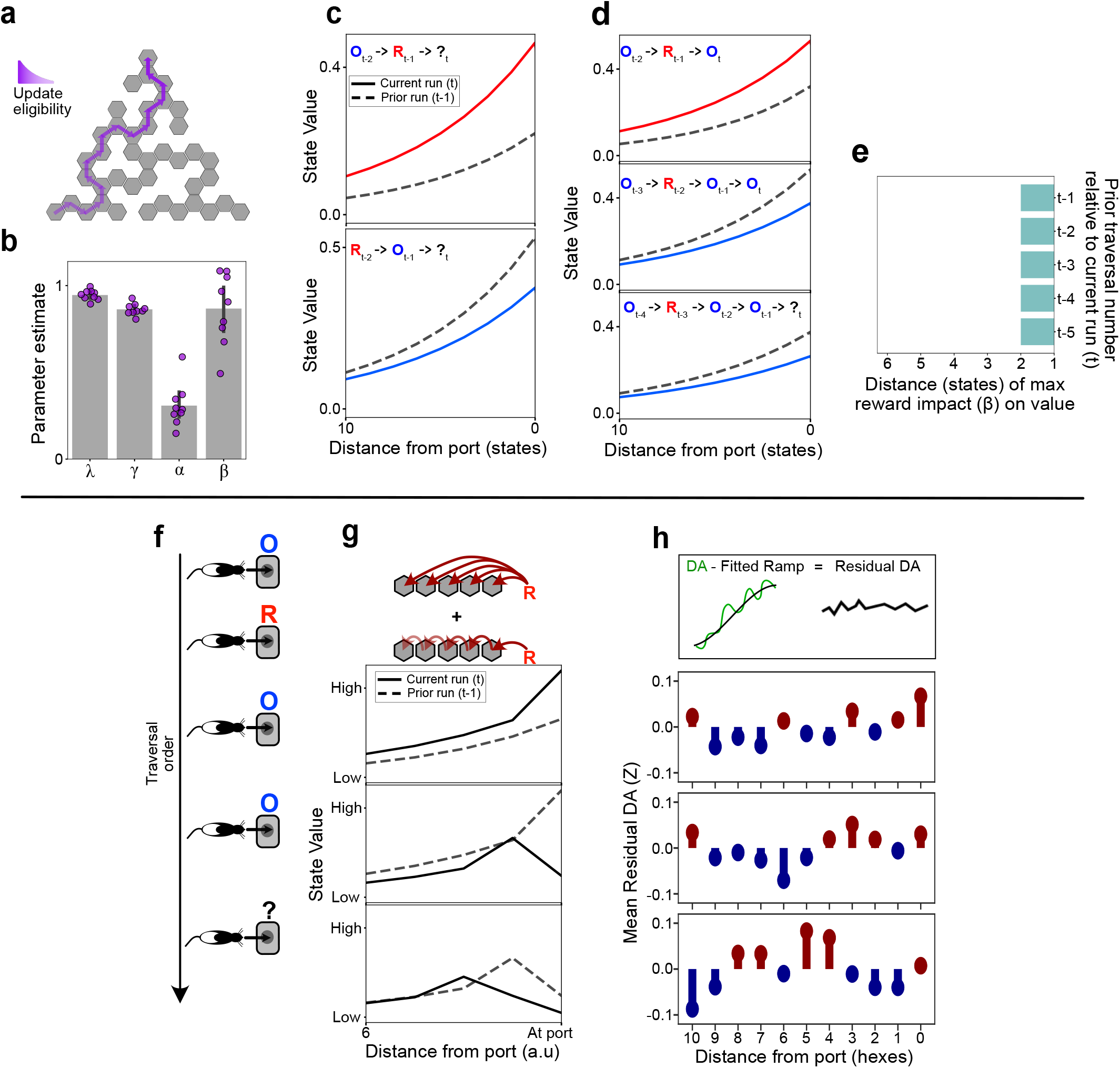
Further comparison of TD algorithms to DA signals. **a**, Schematic of the TD(λ) update algorithm showing the traversed hex-states’ eligibility at the end of a trial. **b**, Estimated parameter values after fitting TD(λ) to each rat’s hex-level DA signal (bars show mean values across rats, dots signify individual-rat estimates, error shows *±* SEM). **c**, Predicted DA value traces based on parameter values from the fitted TD(λ) hex-value learning algorithm. Predicted value function in response to reward and omission, over the same trial sequence as in Fig. 3c. “R” and “O” denote rewards and omissions, respectively, on the t-n previous visits to the port. **d**, Predicted value function after a single reward in a series of omissions, over successive runs of the same path, as in Fig. 5b. **e**, Analyzing the distance from the terminal port in which prior rewards have their strongest impacts (linear regression weights) on state value, as in Fig. 3f-h. Predictions from 1000 simulations of a TD(λ) algorithm, based on parameter values from the fitted TD(λ) hex-value learning algorithm. **f**, Schematic of the traversal sequence from Fig. 5b. **g**, Predicted value function for a linear combination of TD(0) and TD(1) value updates. **h**, Path-wide-update extraction and residual analysis. *Top*, illustration of an individual-trial DA trace, the fitted average ramp for subtraction, and the remaining DA residuals for analysis. *Bottom*, observed mean DA residuals over the three traversals in h.

**Supplementary Figure 5.**
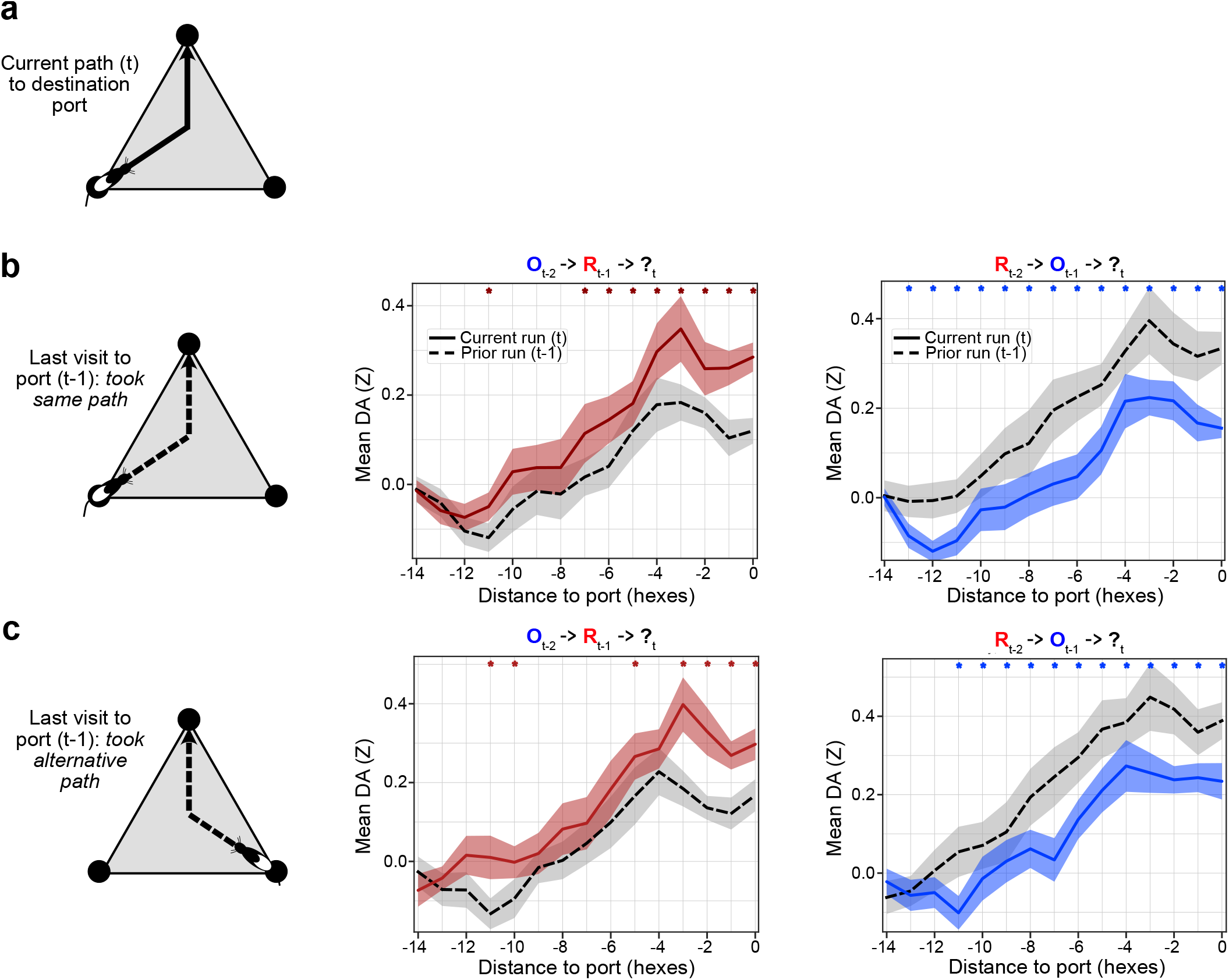
Examining the effects of reward on DA ramping along successive runs to the same port, separated by prior path type. **a**, Schematic illustrating the rat’s current path to the destination port. **b**, *Left*, schematic illustrating trials where the rat previously took the same path to the destination port. *Middle*, mean DA over successive path traversals to the same port, where reward was omitted two visits ago (t-2), but reward was delivered the prior visit (t-1; n=9 rats, 1087 sequences where rat previously took the same path to the destination port). Red asterisks indicate a significant increase in hex-level DA (p < 0.05, one-tailed Wilcoxon signed rank test). “R” and “O” denote rewards and omissions, respectively, on the t-n previous visits to the port. *Right*, same as left but examining the effects of a reward omission on the last visit. Blue asterisks indicate significant DA decrease (p < 0.05, one-tailed Wilcoxon signed rank test; n=9 rats, 1079 sequences where rat previously took the same path to the destination port). **c**, *Left*, schematic illustrating trials where the rat previously took an alternative path to the destination port. *Middle*, same as b middle, but where the rat previously took the alternative path to the destination port (n=9 rats, 592 sequences). *Right*, same as b right, but where the rat previously took the alternative path to the destination port (n=9 rats, 583 sequences).

## Bibliography

1. Richard S Sutton and Andrew G Barto. Reinforcement Learning, second edition: An Introduction. MIT Press, November 2018.

2. Wolfram Schultz, Peter Dayan, and P Read Montague. A neural substrate of prediction and reward. Technical report, 1997.

3. Hannah M Bayer and Paul W Glimcher. Midbrain dopamine neurons encode a quantitative reward prediction error signal. Neuron, 47(1):129–141, 2005.

4. Jeremiah Y Cohen, Sebastian Haesler, Linh Vong, Bradford B Lowell, and Naoshige Uchida. Neuron-type-specific signals for reward and punishment in the ventral tegmental area. Nature, 482, 2012.

5. Ali Mohebi, Jeffrey R Pettibone, Arif A Hamid, Jenny-Marie T Wong, Leah T Vinson, Tommaso Patriarchi, Lin Tian, Robert T Kennedy, and Joshua D Berke. Dissociable dopamine dynamics for learning and motivation. Nature, 570(7759):65–70, June 2019.

6. Andrew S Hart, Robb B Rutledge, Paul W Glimcher, and Paul E M Phillips. Phasic dopamine release in the rat nucleus accumbens symmetrically encodes a reward prediction error term. J. Neurosci., 34(3):698–704, January 2014.

7. Wei-Xing Pan, Robert Schmidt, Jeffery R Wickens, and Brian I Hyland. Dopamine cells respond to predicted events during classical conditioning: evidence for eligibility traces in the reward-learning network. J. Neurosci., 25(26):6235–6242, June 2005.

8. Ryunosuke Amo, Sara Matias, Akihiro Yamanaka, Kenji F Tanaka, Naoshige Uchida, and Mitsuko Watabe-Uchida. A gradual temporal shift of dopamine responses mirrors the progression of temporal difference error in machine learning. Nat. Neurosci., July 2022.

9. Huijeong Jeong, Annie Taylor, Joseph R Floeder, Martin Lohmann, Stefan Mihalas, Brenda Wu, Mingkang Zhou, Dennis A Burke, and Vijay Mohan K Namboodiri. Mesolimbic dopamine release conveys causal associations. Science, page eabq6740, December 2022.

10. Nathaniel D Daw, Yael Niv, and Peter Dayan. Uncertainty-based competition between prefrontal and dorsolateral striatal systems for behavioral control. Nat. Neurosci., 8(12): 1704–1711, December 2005.

11. Yunzhe Liu, Marcelo G Mattar, Timothy E J Behrens, Nathaniel D Daw, and Raymond J Dolan. Experience replay is associated with efficient nonlocal learning. Science, 372(6544), May 2021.

12. Melissa J Sharpe, Hannah M Batchelor, Lauren E Mueller, Chun Yun Chang, Etienne J P Maes, Yael Niv, and Geoffrey Schoenbaum. Dopamine transients do not act as model-free prediction errors during associative learning. Nat. Commun., 11(1):106, January 2020.

13. Brian F Sadacca, Joshua L Jones, and Geoffrey Schoenbaum. Midbrain dopamine neurons compute inferred and cached value prediction errors in a common framework. Elife, 5 (MARCH2016):1–13, 2016.

14. Hiroyuki Nakahara, Hideaki Itoh, Reiko Kawagoe, Yoriko Takikawa, and Okihide Hikosaka. Dopamine neurons can represent Context-Dependent prediction error, 2004.

15. Nathaniel D Daw, Samuel J Gershman, Ben Seymour, Peter Dayan, and Raymond J Dolan. Model-based influences on humans’ choices and striatal prediction errors. Neuron, 69(6): 1204–1215, March 2011.

16. Mitchell F Roitman, Garret D Stuber, Paul E M Phillips, R Mark Wightman, and Regina M Carelli. Dopamine operates as a subsecond modulator of food seeking. J. Neurosci., 24(6): 1265–1271, February 2004.

17. Arif A Hamid, Jeffrey R Pettibone, Omar S Mabrouk, Vaughn L Hetrick, Robert Schmidt, Caitlin M Vander Weele, Robert T Kennedy, Brandon J Aragona, and Joshua D Berke. Mesolimbic dopamine signals the value of work. Nat. Neurosci., 19(1):117–126, January 2016.

18. Anne L Collins, Venuz Y Greenfield, Jeffrey K Bye, Kay E Linker, Alice S Wang, and Kate M Wassum. Dynamic mesolimbic dopamine signaling during action sequence learning and expectation violation. Sci. Rep., 6(October 2015):1–15, 2016.

19. Mark W Howe, Patrick L Tierney, Stefan G Sandberg, Paul E M Phillips, and Ann M Graybiel. Prolonged dopamine signalling in striatum signals proximity and value of distant rewards. Nature, 2013.

20. Kenji Morita and Ayaka Kato. Striatal dopamine ramping may indicate flexible reinforcement learning with forgetting in the cortico-basal ganglia circuits. Front. Neural Circuits, 8(APR): 1–15, 2014.

21. Hyunggoo R Kim, Athar N Malik, John G Mikhael, Pol Bech, Iku Tsutsui-Kimura, Fangmiao Sun, Yajun Zhang, Yulong Li, Mitsuko Watabe-Uchida, Samuel J Gershman, and Naoshige Uchida. A unified framework for dopamine signals across timescales. Cell, 183(6):1600– 1616.e25, December 2020.

22. Akash Guru, Changwoo Seo, Ryan J Post, Durga S Kullakanda, Julia A Schaffer, and Melissa R Warden. Ramping activity in midbrain dopamine neurons signifies the use of a cognitive map. bioRxiv, page 2020.05.21.108886, 2020.

23. Genela Morris, Alon Nevet, David Arkadir, Eilon Vaadia, and Hagai Bergman. Midbrain dopamine neurons encode decisions for future action. Nat. Neurosci., 9(8):1057–1063, 2006.

24. Nathan F Parker, Courtney M Cameron, Joshua P Taliaferro, Junuk Lee, Jung Yoon Choi, Thomas J Davidson, Nathaniel D Daw, and Ilana B Witten. Reward and choice encoding in terminals of midbrain dopamine neurons depends on striatal target. Nat. Neurosci., 19(6): 845–854, June 2016.

25. Vijay Mohan K Namboodiri. How do real animals account for the passage of time during associative learning? Behav. Neurosci., 136(5):383–391, October 2022.

26. D J Foster, R G M Morris, and Peter Dayan. A model of hippocampally dependent navigation, using the temporal difference learning rule. Hippocampus, 10(1):1–16, 2000.

27. Ben Engelhard, Joel Finkelstein, Julia Cox, Weston Fleming, Hee Jae Jang, Sharon Ornelas, Sue Ann Koay, Stephan Y Thiberge, Nathaniel D Daw, David W Tank, and Ilana B Witten. Specialized coding of sensory, motor and cognitive variables in VTA dopamine neurons. Nature, 570(7762):509–513, June 2019.

28. Kazuyuki Samejima, Yasumasa Ueda, Kenji Doya, and Minoru Kimura. Representation of Action-Specific reward values in the striatum. Science, 310(57):1338–1340, 2005.

29. Brian Lau and Paul W Glimcher. Dynamic Response-by-Response models of matching behavior in rhesus monkeys. J. Exp. Anal. Behav., 84(3):555–579, 2005.

30. Nathaniel D Daw, John P O’doherty, Peter Dayan, Ben Seymour, and Raymond J Dolan. Cortical substrates for exploratory decisions in humans. Nature, 2006.

31. Namjung Huh, Suhyun Jo, Hoseok Kim, Jung Hoon Sul, and Min Whan Jung. Model-based reinforcement learning under concurrent schedules of reinforcement in rodents. Learn. Mem., 16(5):315–323, May 2009.

32. Tommaso Patriarchi, Jounhong Ryan Cho, Katharina Merten, Mark W Howe, Aaron Marley, Wei-Hong Xiong, Robert W Folk, Gerard Joey Broussard, Ruqiang Liang, Min Jee Jang, Haining Zhong, Daniel Dombeck, Mark von Zastrow, Axel Nimmerjahn, Viviana Gradinaru, John T Williams, and Lin Tian. Ultrafast neuronal imaging of dopamine dynamics with designed genetically encoded sensors. Science, 2018.

33. Vikram Gadagkar, Pavel A Puzerey, Ruidong Chen, Eliza Baird-Daniel, Alexander R Farhang, and Jesse H Goldberg. Dopamine neurons encode performance error in singing birds. Science, 354(6317):1278–1282, 2016.

34. Yael Niv, Nathaniel D Daw, Daphna Joel, and Peter Dayan. Tonic dopamine: opportunity costs and the control of response vigor. Psychopharmacology, 191(3):507–520, April 2007.

35. Dylan Alexander Simon and Nathaniel D Daw. Neural correlates of forward planning in a spatial decision task in humans. J. Neurosci., 31(14):5526–5533, 2011.

36. Nathaniel D Daw. Trial-by-trial data analysis using computational models. Attention & Performance, XXIII, 2009.

37. J C Horvitz. Mesolimbocortical and nigrostriatal dopamine responses to salient non-reward events. Neuroscience, 96(4):651–656, 2000.

38. Ethan S Bromberg-Martin, Masayuki Matsumoto, and Okihide Hikosaka. Dopamine in motivational control: Rewarding, aversive, and alerting, December 2010.

39. Peter Redgrave and Kevin Gurney. The short-latency dopamine signal: a role in discovering novel actions? Nat. Rev. Neurosci., 7(12):967–975, December 2006.

40. Matthew P H Gardner, Geoffrey Schoenbaum, and Samuel J Gershman. Rethinking dopamine as generalized prediction error, 2018.

41. Emilie C J Syed, Laura L Grima, Peter J Magill, Rafal Bogacz, Peter Brown, and Mark E Walton. Action initiation shapes mesolimbic dopamine encoding of future rewards. Nat. Neurosci., 19(1):34–36, January 2016.

42. Mayank Agrawal, Marcelo G Mattar, Jonathan D Cohen, and Nathaniel D Daw. The temporal dynamics of opportunity costs: A normative account of cognitive fatigue and boredom. Psychol. Rev., 129(3):564–585, April 2022.

43. Ian Osband, Charles Blundell, Alexander Pritzel, and Benjamin Van Roy. Deep exploration via bootstrapped DQN. February 2016.

44. Mark E Walton and Sebastien Bouret. What is the relationship between dopamine and effort? Trends Neurosci., pages 1–13, 2018.

45. John D Salamone, Michael S Cousins, and Sherri Bucher. Anhedonia or anergia? effects of haloperidol and nucleus accumbens dopamine depletion on instrumental response selection in a t-maze cost/benefit procedure. Technical report, 1994.

46. M S Cousins, A Atherton, L Turner, and J D Salamone. Nucleus accumbens dopamine depletions alter relative response allocation in a t-maze cost/benefit task. Behav. Brain Res., 1996.

47. Shunsuke Kobayashi and Wolfram Schultz. Influence of reward delays on responses of dopamine neurons. J. Neurosci., 28(31):7837–7846, July 2008.

48. Wei Wei, Ali Mohebi, and Joshua D Berke. A spectrum of time horizons for dopamine signals. May 2022.

49. Samuel J Gershman, Ahmed A Moustafa, and Elliot A Ludvig. Time representation in reinforcement learning models of the basal ganglia. Front. Comput. Neurosci., 7(January): 1–8, 2014.

50. John G Mikhael, Hyunggoo R Kim, Naoshige Uchida, and Samuel J Gershman. The role of state uncertainty in the dynamics of dopamine. Curr. Biol., 32(5):1077–1087.e9, March 2022.

51. David J Foster and Matthew A Wilson. Reverse replay of behavioural sequences in hippocampal place cells during the awake state. Nature, 440(30), 2006.

52. Annabelle C Singer and Loren M Frank. Rewarded outcomes enhance reactivation of experience in the hippocampus. Neuron, 64(6):910–921, December 2009.

53. R Ellen Ambrose, Brad E Pfeiffer, and David J Foster. Reverse replay of hippocampal place cells is uniquely modulated by changing reward. Neuron, 91, 2016.

54. Helen C Barron, Hayley M Reeve, Renée S Koolschijn, Pavel V Perestenko, Anna Shpektor, Hamed Nili, Roman Rothaermel, Natalia Campo-Urriza, Jill X O’Reilly, David M Bannerman, Timothy E J Behrens, and David Dupret. Neuronal computation underlying inferential reasoning in humans and mice. Cell, 183(1):228–243.e21, October 2020.

55. Baburam Bhattarai, Jong Won Lee, and Min Whan Jung. Distinct effects of reward and navigation history on hippocampal forward and reverse replays. Proc. Natl. Acad. Sci. U. S. A., 117(1):689–697, January 2020.

56. Marcelo G Mattar and Nathaniel D Daw. Prioritized memory access explains planning and hippocampal replay. Nat. Neurosci., 21(11):1609–1617, November 2018.

57. Hado van Hasselt, Sephora Madjiheurem, Matteo Hessel, David Silver, André Barreto, and Diana Borsa. Expected eligibility traces. AAAI, 35(11):9997–10005, May 2021.

58. Anna Harutyunyan, Will Dabney, Thomas Mesnard, Mohammad Azar, Bilal Piot, Nicolas Heess, Hado van Hasselt, Greg Wayne, Satinder Singh, Doina Precup, and Remi Munos. Hindsight credit assignment. December 2019.

59. David Silver, Aja Huang, Chris J Maddison, Arthur Guez, Laurent Sifre, George Van Den Driessche, Julian Schrittwieser, Ioannis Antonoglou, Veda Panneershelvam, Marc Lanctot, Sander Dieleman, Dominik Grewe, John Nham, Nal Kalchbrenner, Ilya Sutskever, Timothy Lillicrap, Madeleine Leach, Koray Kavukcuoglu, Thore Graepel, and Demis Hassabis. Mastering the game of go with deep neural networks and tree search. Nature, 2016.

60. Brad E Pfeiffer and David J Foster. Hippocampal place-cell sequences depict future paths to remembered goals. Nature, 497(7447):74–79, May 2013.

61. Andrew M Wikenheiser and David Redish. Hippocampal theta sequences reflect current goals. Nat. Neurosci., 18(2), 2015.

62. Kenneth Kay, Jason E Chung, Marielena Sosa, Jonathan S Schor, Mattias P Karlsson, Margaret C Larkin, Daniel F Liu, and Loren M Frank. Constant sub-second cycling between representations of possible futures in the hippocampus. Cell, 180(3):552–567.e25, February 2020.

63. Alison E Comrie, Loren M Frank, and Kenneth Kay. Imagination as a fundamental function of the hippocampus. Philos. Trans. R. Soc. Lond. B Biol. Sci., 377(1866):20210336, December 2022.

64. Adam Johnson and A David Redish. Neural ensembles in CA3 transiently encode paths forward of the animal at a decision point. J. Neurosci., 2007.

65. S M Nicola. The flexible approach hypothesis: Unification of effort and Cue-Responding hypotheses for the role of nucleus accumbens dopamine in the activation of Reward-Seeking behavior. Journal of Neuroscience, 2010.

66. S Ikemoto and J Panksepp. The role of nucleus accumbens dopamine in motivated behavior: a unifying interpretation with special reference to reward-seeking. Brain Res. Brain Res. Rev., 31(1):6–41, December 1999.

67. Tanmay Nath, Alexander Mathis, An Chi Chen, Amir Patel, Matthias Bethge, and Mackenzie Weygandt Mathis. Using DeepLabCut for 3D markerless pose estimation across species and behaviors. Nat. Protoc., 14(7):2152–2176, 2019.

68. Ekaterina Martianova, Sage Aronson, and Christophe D Proulx. Multi-Fiber photometry to record neural activity in Freely-Moving animals. J. Vis. Exp., (152):1–9, October 2019.

69. Silviu Pitis. Source traces for temporal difference learning. AAAI, 32(1), April 2018.

70. Quentin J M Huys, Roshan Cools, Martin Gölzer, Eva Friedel, Andreas Heinz, Raymond J Dolan, and Peter Dayan. Disentangling the roles of approach, activation and valence in instrumental and pavlovian responding. PLoS Comput. Biol., 7(4):e1002028, April 2011.

71. D Oakes. Direct calculation of the information matrix via the EM. J. R. Stat. Soc. Series B Stat. Methodol., 61(2):479–482, April 1999.

